# CDK4/6 inhibition and dsRNA sensor agonism co-operate to enhance anti-cancer effects through ER stress and immune modulation of tumour cells

**DOI:** 10.1101/2022.09.28.508679

**Authors:** Victoria Roulstone, Joan Kyula, James Wright, Lu Yu, Aida Barreiro Alonso, Miriam Melake, Jyoti Choudhary, Richard Elliott, Christopher J. Lord, David Mansfield, Nik Matthews, Ritika Chauhan, Victoria Jennings, Charleen Chan, Holly Baldock, Francesca Butera, Elizabeth Appleton, Pablo Nenclares, Malin Pederson, Shane Foo, Emmanuel C. Patin, Antonio Rullan, Tencho Tenev, Pascal Meier, Jacob Van Vloten, Richard Vile, Hardev Pandha, Alan Melcher, Martin McLaughlin, Kevin Harrington

**Affiliations:** The Institute of Cancer Research, London, UK; The CRUK Gene Function Laboratory and Breast Cancer Now Toby Robins Research Centre, The Institute of Cancer Research, London, UK; Mayo Clinic, Rochester, Minnesota, US; University of Surrey, Guildford, UK

**Keywords:** CDK4/6 inhibitor, palbociclib, unfolded protein response (UPR), integrated stress response, ER stress, oncolytic virus, reovirus type 3 Dearing, RNA sensing, nucleic acid sensing, RIG-I, MDA5, TLR3, interferon

## Abstract

Cytoplasmic pattern recognition receptors (PRRs) for double-stranded RNA (RIG-I/MDA5) are key mediators of anti-viral responses. PRR agonists, such as dsRNA oncolytic Reovirus type 3 Dearing (Rt3D), potently activate RNA sensors. We used an unbiased cytotoxicity screen to reveal synergistic drug-virotherapy combinations and found potent effects of Rt3D combined with the CDK4/6 inhibitor, palbociclib. The combination augmented oncolytic virus-induced endoplasmic reticulum (ER) stress/unfolded protein response (UPR) and the expression and activation/signaling of RNA sensors. Combined Rt3D-palbociclib treatment potently increased interferon production and signaling, and knockdown studies implicated key UPR proteins and the RNA sensor, RIG-I, as essential to the phenotype observed. Further experiments, using canonical RIG-I agonists and an ER stress inducer, thapsigargin, confirmed cross-talk between RNA sensing and ER stress pathways that augmented cancer cell death and interferon production. Combined Rt3D-palbociclib also increased innate immune activation within tumour cells and IFN-induced HLA expression. Analysis of the immunopeptidome revealed changes to HLA-captured peptides with Rt3D-palbociclib, including altered expression of peptides from cancer/testis antigens (CTA) and endogenous retroviral elements (ERVs). Our findings highlight cross-talk between UPR signaling and RNA-mediated PRR activation as a means of enhancing anti-cancer efficacy with potential pro-immunogenic consequences. This has implications for future clinical development of PRR agonists and oncolytic viruses, and broadens the therapeutic remit of CDK4/6 inhibitors to include roles as both ER stress and dsRNA PRR sensitizers.

## Palbociclib enhances anti-cancer efficacy of reovirus Type 3 Dearing (Rt3D) and the viral defense response

We performed a high-throughput small molecule screen using a non-biased approach to seek synergistic drug-virus interactions. The A375 BRAF^V600E^-mutant melanoma cell line was selected for screening due to the susceptibility of melanoma to Rt3D infection^1^. Compounds used are shown (**Fig. S1A**). The CDK4/6 inhibitor (CDK4/6i), palbociclib, enhanced cytotoxicity across a range of viral doses (**Fig. 1A, S1B**). Palbociclib-mediated CDK4/6 inhibition repressed phospho-retinoblastoma protein (Rb) in *RB*-wild-type cells inducing G1/S cell cycle arrest ^2^. This was corroborated in A375 and other melanoma cell lines (**Fig. S1C**). To validate findings from the screen, the Rt3D-palbociclib combination was tested with orthogonal methods in A375 and other melanoma, head and neck and breast cancer cells, including those with acquired palbociclib resistance (**Fig. 1B, C, S1D-I**). The requirement for intact Rb signaling was shown in A2058 *RB*-null cells, in which palbociclib failed to mediate G1 arrest or augment Rt3D-induced cytotoxicity (**Fig. 1D, S1J**). Efficacy of Rt3D-palbociclib combination therapy was tested *in vivo* using human A375 tumour-bearing CD1 nude mice and in an immunocompetent model bearing murine 4434 BRAF-mutant melanoma (**Fig. 1E**).

**Figure 1.**
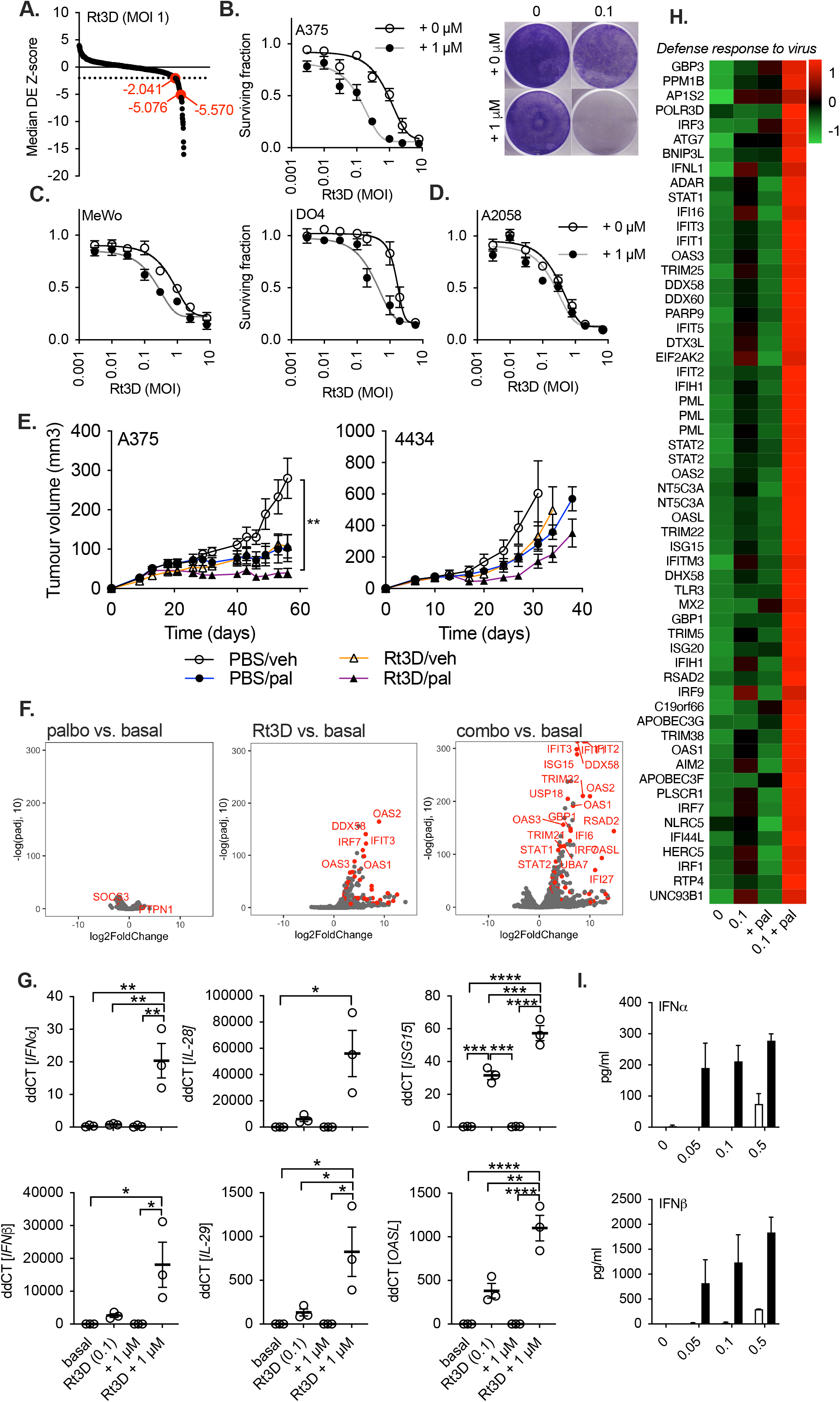
Palbociclib enhances Rt3D cytotoxicity and virus defence response. **A**, A375 were treated with Rt3D (MOI 1) and 80 compounds at various doses. Low Z scores (below -2, dashed line) indicate sensitisers (palbociclib is shown in red, with relative Z scores that represent PD-0332991 doses 1μM (-2.041) and PF-332991 doses at 1μM (-5.076) and 0.5μM (-5.570)). **B**, Cell viability in A375 cells treated with Rt3D +/-palbociclib (1 μM) by MTT (left) and crystal violet (right) 72 hours after treatment (± SEM, *n* = 3). **C**, Cell viability in melanoma cells MeWo (BRAF/RAS wild-type), and DO4 (Ras mutant) treated with Rt3D +/-palbociclib (1 μM) by MTT (72 hours) (± SEM, *n* = 3). **D**, Cell survival is not altered in *RB*-null A2058 cells treated with Rt3D +/-palbociclib (1 μM) shown by MTT, 72 hours (± SEM, *n* = 3). **E**, Tumour volumes for animals bearing A375 tumours treated with an intratumoural injection of Rt3D (1 × 10^6^ pfu) +/-palbociclib/vehicle (left), tumour volumes for C57BL/6 mice bearing murine BRAF mutant 4434 tumours treated with an intratumoural injection of Rt3D (1 × 10^6^ pfu) -/+ palbociclib (right). **F**, Volcano plot of RNA sequencing showing upregulated and downregulated genes for A375 cells treated with palbociclib (1 μM), Rt3D (MOI 0.1) or the combination compared to basal at 48 hours. Highlighted genes in red belong to the GSEA set ‘reactome interferon signalling’. **G**, RT-qPCR of *IFN*α, *IFN*β, *IL-28, IL-29, ISG15* and *OASL* in A375 cells treated with Rt3D +/-palbociclib (1 μM) at 48 hours (± SEM, *n* = 3). **H**. Proteomic analysis of A375 cells treated with Rt3D (0.1) in combination with palbociclib (1 μM) reveal upregulated (red) and downregulated (green) proteins categorised under the GO term ‘defense reponse to virus’ (24 hours). **I**. IFNα and IFNβ, measured in cell-free supernatant from Rt3D-treated samples (MOI 0.05-0.5) +/-palbociclib (1 μM) by ELISA.

To probe mechanistic interactions between Rt3D and palbociclib, we analysed the transcriptome and the proteome. At the transcriptomic level, data indicated a more pronounced interferon signature in tumour cells with combination treatment (**Fig. 1F**). These data were corroborated in separate RT-qPCR experiments for type I *IFN*α, *IFN*β, type III *IL-28* and *IL-29* and IFN-stimulated genes *ISG15* and *OASL* (**Fig. 1G**). Increased mouse *IFN*α, was observed for the mouse melanoma cell line (4434) with Rt3D-palbociclib combination therapy (**Fig. S1K**), despite the absence of increased cell kill with the combination *in vitro* (**Fig. S1L**). At the proteomic level, proteins detected within the GO term ‘defence response to virus’ were upregulated in the Rt3D-palbociclib combination treatment (**Fig. 1H**). Finally, to confirm that IFN expression translated to IFN protein, an ELISA was performed for IFNα/β and IL28/29 (**Fig. 1I, Fig. S1M**).

### Rt3D-palbociclib-induced interferon signaling increases antigen processing machinery and immunogenicity

Proteomic analyses of proteins with GO terms ‘immune response’, ‘interferon regulation/response’ and ‘antigen-processing and presentation’ showed increases in the expression of class I and II HLA-related proteins with Rt3D-palbociclib, that were also evident in transcriptomic analysis (**Fig. 2A, Fig. S2A-C**). FACS analysis of non-permeabilized cells confirmed cell surface expression of HLA class-I and class-II (**Fig. 2B, S2D**), which was abrogated by ruxolitinib-mediated inhibition of IFN signaling (**Fig. 2C, S2E**). Analysis of immunoproteasome subunits showed that Rt3D induced a switch from the standard proteasome to the immunoproteasome, as well as changes in transporter associated with antigen processing (TAP) and ERAP proteins (**Fig. 2D, S2F**). Given these change, and the fact that Rt3D-palbocilib increases cell surface class I HLA expression, we performed immunopeptidomic analyses to look for evidence of treatment-related modulation of peptide presentation. We focused on potential tumour-associated antigens and showed changes in presentation of multiple cancer/testis antigens (CTA) with Rt3D-palbociclib treatment (**Fig. 2E**). A total of 63 CTA immunopeptides from 44 genes were identified, many showing increased expression upon combination treatment. In particular, Rt3D-palbociclib increased the expression of CTA derived from the protein MAGEA4. MAGEA4 mRNA expression levels were not altered by Rt3D-palbociclib treatment, nor by disruption to IFN signalling by the JAK/STAT inhibitor, ruxilitinib. However, protein levels dropped with treatment, suggesting increased turnover by the proteasome (**Fig. 2F**).

**Figure 2.**
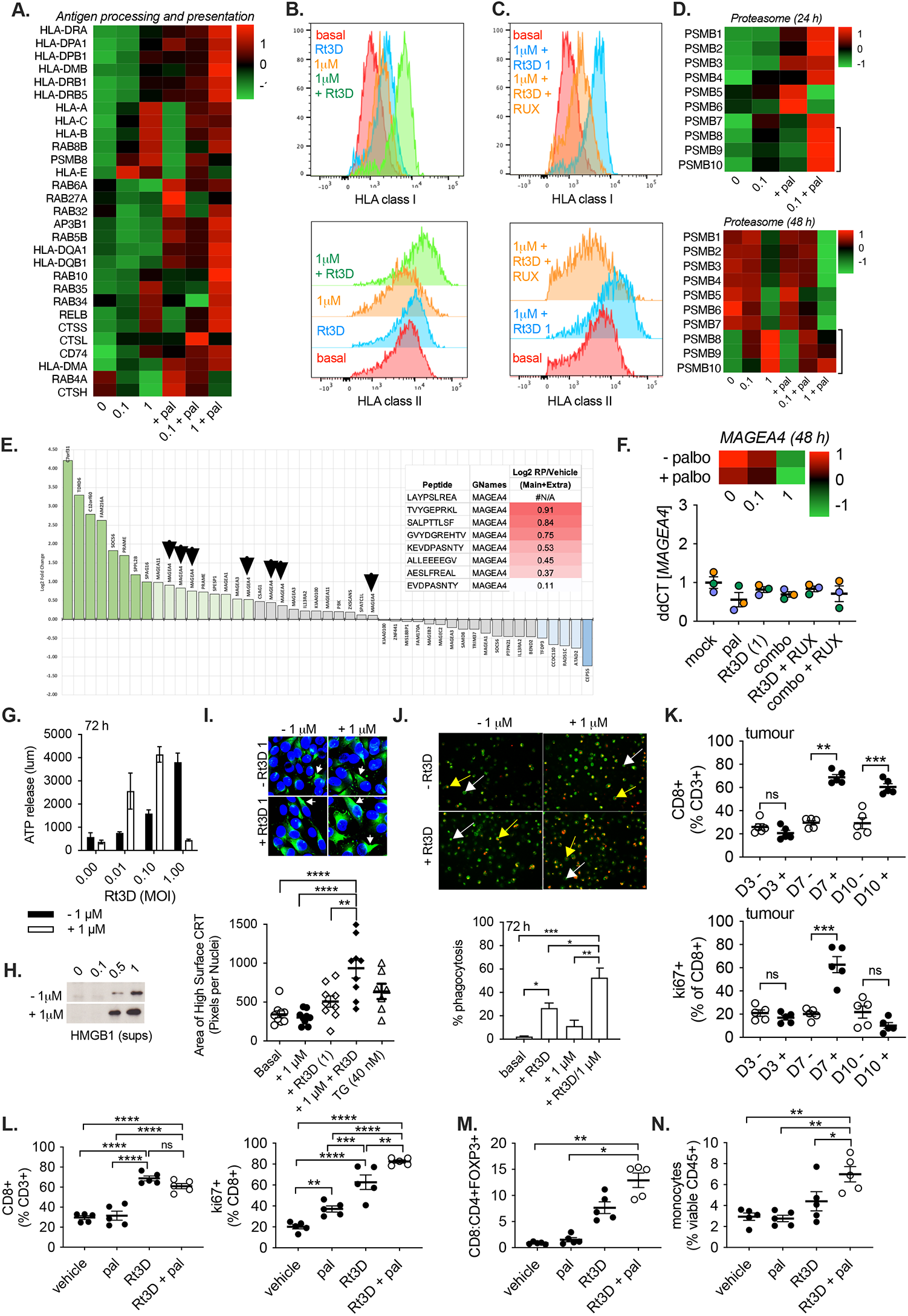
Rt3D-palbociclib increases antigen processing machinery and immunogenicity. Proteomic analysis of A375 cells treated with Rt3D (0.1-1) +/-palbociclib (1 μM) reveal upregulated (red) and downregulated (green) proteins categorised under the GO term ‘antigen processing and presentation’ (48 hours). **B**. Expression of HLA-A,B,C (class I) and HLA-DR,DP,DQ (class II) in A375 cells treated with Rt3D +/-palbociclib by FACS analysis. **C**, Expression of HLA-A,B,C (class I) and HLA-DR,DP,DQ (class II) in A375 cells treated with Rt3D plus palbociclib +/-the JAK/STAT inhibitor, ruxilitinib (RUX). **D**. Proteomic analysis of A375 cells treated with Rt3D (0.1-1) +/-palbociclib (1 μM) reveal upregulated (red) and downregulated (green) proteosome subunits (24 hours, above, 48 hours, below). PSMB8, 9 and 10 (highlighted), are incorporated into the proteosome to switch the standard proteasome into an immunoproteasome. **E**. Fold-change in CTA petides from HLA-I with Rt3D-palbociclib versus basal conditions. Peptides belonging to the MAGEA4 protein are highlighted (1-8). **F**. Proteomic analysis of A375 cells treated with Rt3D (0.1-1) +/-palbociclib (1 μM) reveal upregulated (red) and downregulated (green) MAGEA4 protein (48 hours, top). RT-qPCR of *MAGEA4* in A375 cells treated with Rt3D +/-palbociclib, -/+ ruxilitinib (RUX) (1 μM) at (48 hours, below). **G**. ATP release by cell titre glo assay in supernatants from A375 cells treated with indicated Rt3D doses -/+ palbociclib (1 μM) (± SEM, *n* = 3). **H**, HMGB1 release by western blot of supernatants of treated A375 cells. **I**, Calreticulin (CRT, white arrows) cell surface expression by confocal microscopy of treated A375 cells, quantified for 3 biological repeats (below). **J**, Pictomicrographs of macrophages stained with CD11b-FITC co-cultured with treated, phrodo-stained tumour cells (red), showing engulfed (yellow arrows) and non-engulfed (white arrows) cells. Percentages of macrophage population that dual-stain for phrodo, for 3 biological repeats are shown below. **K**. C57BL/6 mice bearing MOC1 tumours were harvested 3 days (D3), 7 days (D7) or 10 days (D10) after a single injection of 5 × 10^6^ pfu Rt3D (+) or sham injection (-), and stained with extra- and intra-cellular antibodies to profile the immune infiltrate. Data show proportions (through percentage analysis) of CD3+ cells gated from viable cells. Data show the percentage CD8+ cells from the viable CD3+ population (above), and intracellular stained ki67+ are shown as a percentage of the viable cells that stain for CD3+ and CD8+. Unpaired t tests were used to compare populations between Rt3D treated vs. PBS sham for each time point, n=6 animals per group. **L**. C57BL/6 mice bearing MOC1 tumours were treated with palbociclib (100 mg/kg daily by oral gavage). After 3 doses of palbociclib, tumours received a single injection of 5 × 10^6^ pfu Rt3D. Tumours were harvested at day 7 after injection and weighed before staining with extra- and intra-cellular antibodies to profile the immune infiltrate. From the viable CD3+ population, percentage CD8+ are shown (left), from within the CD3+ CD8+ population and intracellular stained ki67+ CD8+ cells are shown as a percentage of CD8+ cells (right). **M**, T cell ratios were calculated for CD8+:CD4+ FOXP3+ for each treatment arm. **N**, Data show proportions (through percentage analysis) of CD45+CD11b+Ly6C+ cells (monocytes) for each treatment arm. A one-way ANOVA was used with the p value corrected for multiple comparisons against the Rt3D + pal group, n=6 animals per group.

Next, we looked for Rt3D-palbociclib-induced tumour cell-intrinsic adjuvanticity through immunogenic cell death (ICD). ICD has been previously described for single-agent CDK4/6 inhibition^3,4^. Surrogate ICD markers (ATP and HMGB1 release, cell-surface calreticulin) were increased following combined Rt3D-palbociclib treatment relative to either agent alone (**Fig. 2G-I**). Given these observations of increased antigen processing/presentation and ICD, which serve as ‘find me’ and ‘eat me’ signals^5^, we next tested if this resulted in elevated phagocytosis by cultured immune cells. CD11b-FITC-labelled PBMC-derived human macrophages were co-cultured with tumour cells that had prior staining with pHrodo, a dye that fluoresces in acidic environments, such as phagolysosomes^6^. Macrophage engulfment of Rt3D-infected tumour cells was significantly enhanced by palbociclib, measured by dual-staining (**Fig. 2J, S2G**). Taken together, these results support the notion that Rt3D-palbociclib treatment of tumour cells may increase activation of both innate and adaptive anti-tumour immunity.

To develop this idea, we next investigated the infiltration of T cells and myeloid cells following Rt3D-palbociclib treatment in an immunocompetent model *in vivo*. Analysis of tumour-infiltrating CD8+ cells on day 3, 7 and 10 after a single intratumoural injection of Rt3D into MOC1 tumours, revealed an increase in CD8+ T cells by day 7, that persisted on day 10, with increased staining for the proliferation marker ki67 only on day 7 (**Fig. 2K**). On day 7, the Rt3D-palbociclib combination therapy did not increase the proportion of CD8+ T cells over that seen with Rt3D treatment alone, but did boost ki67+CD8+ T cells (**Fig. 2L**). Furthermore, Rt3D-palbociclib therapy increased the CD8:T reg ratio over the single agent counterparts (**Fig. 2M, S2H**). In the myeloid compartments, Rt3D-palbociclib increased the infiltration of monocytes within the TME (**Fig. 2N**), although there was a reduction in DCs and macrophages with palbociclib or Rt3D-palbociclib (**Fig. S2I**), potentially representing relocation of these antigen presenting cells to draining lymph nodes, or a loss of these immune cell subtypes. Rt3D-palbociclib treatment also resulted in elevated levels of PD-L1 expression on monocytes and macrophages (**Fig. S2J**). Taken together, Rt3D-palbociclib treatment increases the infiltration of proliferative CD8+ T cells and the CD8:T reg ratio, and alters the proportions of myeloid cells and their expression of PD-L1 within the TME, which may warrant assessment of Rt3D-palbociclib in additional combination with anti-PD-1 therapy in future, more primarily immune-focused studies.

### Rt3D-palbociclib increases RNA sensor expression and JAK/STAT dependent expression of ERV species

To determine the palbociclib-specific effects within combination therapy, transcriptomic analysis of Rt3D-palbociclib was compared with Rt3D. This revealed marked downregulation in genes sets associated with chromatin accessibility and nucleosome organization. Histone mRNA expression was significantly downregulated, a result that was corroborated in all histones detected in proteomic analysis (**Fig. 3A, B**). Additionally, analysis of proteins with the GO term ‘DNA modification’ revealed decreases in DNA methyltransferases, including suppression of DNA methyltransferase-1 (DNMT1) and EZH2, regardless of the presence of Rt3D (**Fig. 3C**). It has been reported that reduced DNMT function, either through CDK4/6 inhibition or direct DNMT-inhibition, can trigger release of endogenous retroviral elements (ERV) RNA species, which can act as tumour-associated antigens and stimulate interferon responses^7,8^. The RNA sensors RIG-I (*DDX58*), MDA-5 (*IFIH1*) and *TLR3* were all significantly upregulated with Rt3D-palbociclib relative to single-agent counterparts (**Fig. 3D**). To determine the contribution of RNA sensors to combined therapeutic efficacy, knockdown of RIG-I (but not of MDA-5 or TLR3) rescued cells from the cytotoxicity of Rt3D-palbociclib combination (**Fig. 3E, Fig. S3A**). RIG-I knockdown also reduced the expression of virus-induced IFN response genes (**Fig. S3B**), implicating its importance both in cell death and the host response to viral infection.

**Figure 3.**
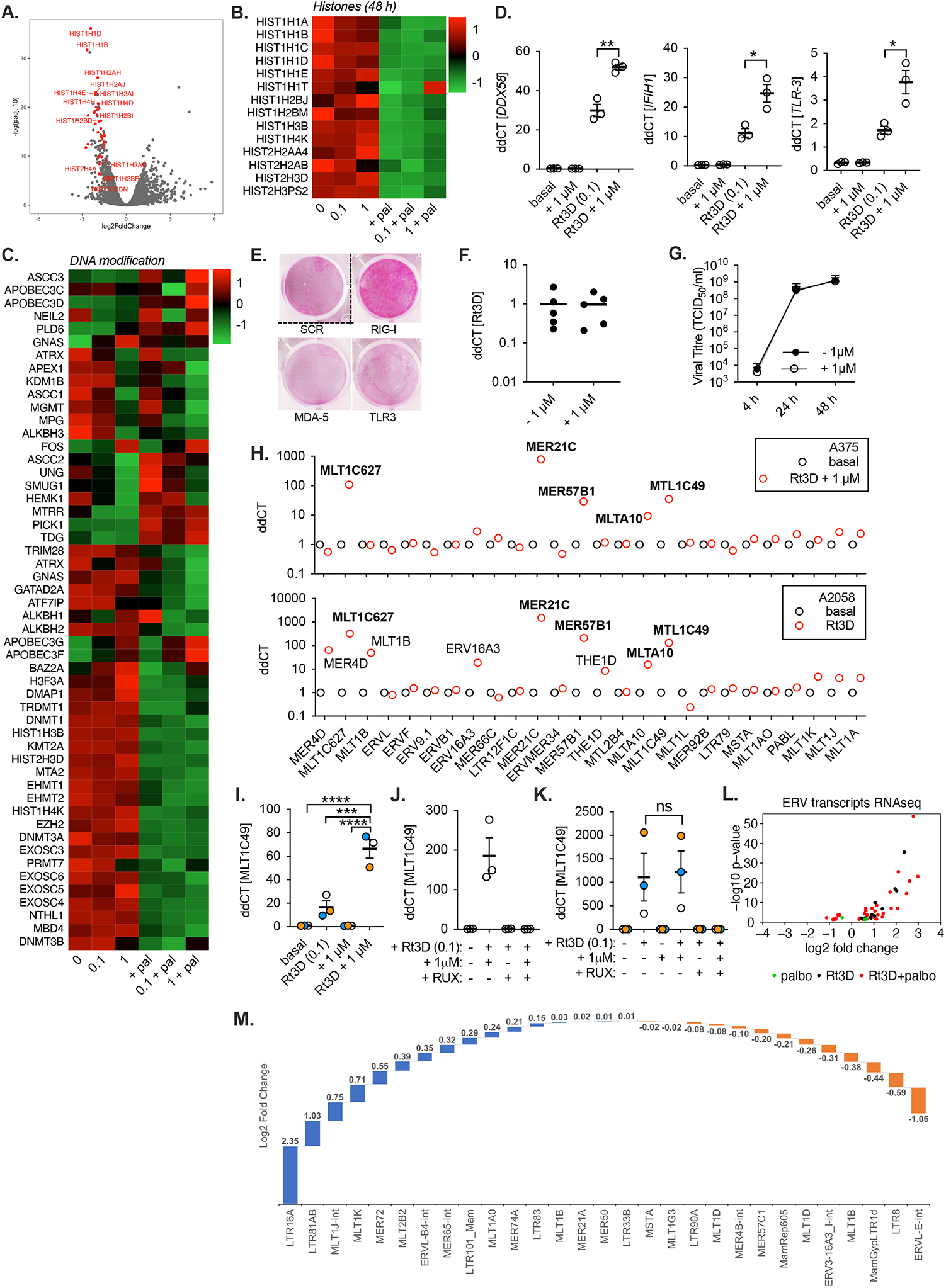
Rt3D-palbociclib increases RNA sensor expression and RNA ERV species expression that is dependent on JAK/STAT signalling. ERV peptides are found in the immunoproteome. **A**. Volcano plot of RNA sequencing showing upregulated and downregulated genes for A375 cells treated with Rt3D (MOI 0.1) plus palbociclib (1 μM), compared to Rt3D alone, at 48 hours. **B**. Proteomic analysis of A375 cells treated with Rt3D (0.1-1) +/-palbociclib (1 μM) reveal upregulated (red) and downregulated (green) histone proteins (48 hours). **C**. Proteomic analysis of A375 cells treated with Rt3D (0.1-1) +/-palbociclib (1 μM) categorised under the GO term ‘DNA modification’ (48 hours). **D**. RT-qPCR of RIG-I (*DDX58*), MDA-5 (*IFIH1*) and *TLR-3* in A375 cells treated with Rt3D plus palbociclib (1 μM) at 48 hours (± SEM, *n* = 3). **E**. siRNA silencing of RNA sensors in A375 cells treated with Rt3D (0.1) -/+ palbociclib (1 μM) by SRB. Data are representative of 3 independent repeats. **F**. RT-qPCR of Rt3D genome in A375-infected cells +/- palbociclib (1 μM) at 48 hours. **G**, Replication of Rt3D by one-step virus growth assay for A375 +/-palbociclib (1 μM). **H**. RT-qPCR for the expression of a panel of 26 previously described ERVs in A375 (above) or A2058 (below), treated with combination therapy, or Rt3D respectively. **I**. RT-qPCR of *MLT1C49* in A375 cells treated with Rt3D +/-palbociclib (1 μM) at 48 hours (± SEM, *n* = 3). **J**. *MLT1C49* expression in A375 cells treated with Rt3D and/or palbociclib (1 μM) +/-JAK/STAT inhibitor (ruxolitinib, RUX) at 48 hours (± SEM, *n* = 3). **K**. RT-qPCR of *MLT1C49* in *RB-*null A2058 cells treated with Rt3D +/-palbociclib (1 μM), +/-ruxolitinib (RUX), at 48 hours (± SEM, *n* = 3). **L**. Volcano plot of RNA sequencing data showing upregulated and downregulated ERVs in A375 cells treated with Rt3D -/+ palbociclib compared to basal at 48 hours (using TETool kit). **M**. Fold-change in ERV peptides from captured HLA-I with Rt3D-palbociclib versus basal conditions. ERV Peptides Filtered: 5% Peptide FDR, Confirmed TMT Quant, Single Genomic Loci, Predicted as MHC Binding.

As Rt3D is a dsRNA virus, we surmised that enhanced interferon signaling could be driven by increased Rt3D genome levels. However, RT-qPCR analysis of Rt3D RNA showed that palbociclib did not alter viral RNA levels (**Fig. 3F**), nor did it increase viral replication (**Fig. 3G**). Therefore, we investigated the possibility that expression of ERV RNA species is detected by the high levels of RNA sensors, contributing to the increased IFN response. A375 cells treated with Rt3D-palbociclib were screened for 26 previously described ERVs. Treatment induced expression of 5 ERVs - *MLT1C49, MER21C, MLTA10, MLT1C627* and *MER57B1*. Similar testing in the *RB*-null cell line, A2058, revealed that Rt3D also induced expression of the same 5 ERVs, but additionally *MER4A, MLT1B, ERV16A3* and *THE1D* (**Fig. 3H**). Palbociclib increased the expression of 3 of 5 Rt3D-induced ERVs (*MLT1C49, MER21C, MLTA10*) in A375 (**Fig. 3I, Fig. S3C**). However, palbociclib had no effect on these Rt3D-induced ERVs in the *RB*-null A2058 cell line (**Fig. 3K, Fig. S3E**). Since the expression of ERVs was dependent on Rt3D, but not palbociclib, we considered that their expression was likely IFN-dependent, which was consistent with the location of the 5 ERVs expressed in both cell lines within STAT1-associated genes (**Table S1**). Abrogation of ERV expression with ruxolitinib confirmed their expression was dependent on interferon/JAK/STAT signaling (**Fig. 3J, K, Fig. S3D, E**). RNA sequencing corroborated that Rt3D and Rt3D-palbociclib induced ERV expression (**Fig 3L, Fig. S3F**). Taken together, these data show that palbociclib drives marked alterations to chromatin, including reduced expression of histones and DNA methyltransferases. Rt3D treatment drives expression of ERVs, which are dependent on JAK/STAT signalling. Interestingly, we observed neither ERV transcription nor ISG induction due to single-agent palbociclib. Rather, palbociclib enhances RNA sensor expression and transcription of select ERVs induced by Rt3D. Finally, we analysed the immunopeptidome, and found that ERV elements can be presented on HLA class I molecules, and that the expression of these is altered by Rt3D-palbociclib treatment (**Fig. 3M**). Analysis of the RNA expression levels of the ERV sequences encoding epitopes found on the immunopeptidome revealed no correlation with treatment (**Fig S3G, H**), highlighting the likely complexity of the roles of differing ERVs in triggering an IFN response or being translated, processed and presented as potential tumour-associated antigens.

### Rt3D-palbociclib augments ER stress

Since palbociclib exerts a G1 cell cycle arrest, we explored the cell cycle effects with combination therapy. Palbociclib induced an increase in the G1 compartment, as expected. However, the combination of Rt3D-palbociclib led to a significantly increased sub-G1 population (**Fig. 4A**). Compared with single-agent therapies, combined Rt3D-palbociclib did not alter levels of phosho-Rb, CDK4, Cyclin D1, E1 or A2 at the protein level (**Fig. S4A**). Conversely, western analysis confirmed that Rt3D-induced caspase-3 and PARP cleavage was enhanced by palbociclib (**Fig. 4B**). Mechanistically, this suggested an apoptotic event, and we confirmed, using a panel of caspase inhibitors, that the caspase-4 inhibitor, Z-YVAD-FMK, or siRNA-mediated caspase-4 knockdown rescued the phenotype of the Rt3D-palbociclib combination treatment (**Fig. 4C, D, Fig. S4B, C**).

**Figure 4.**
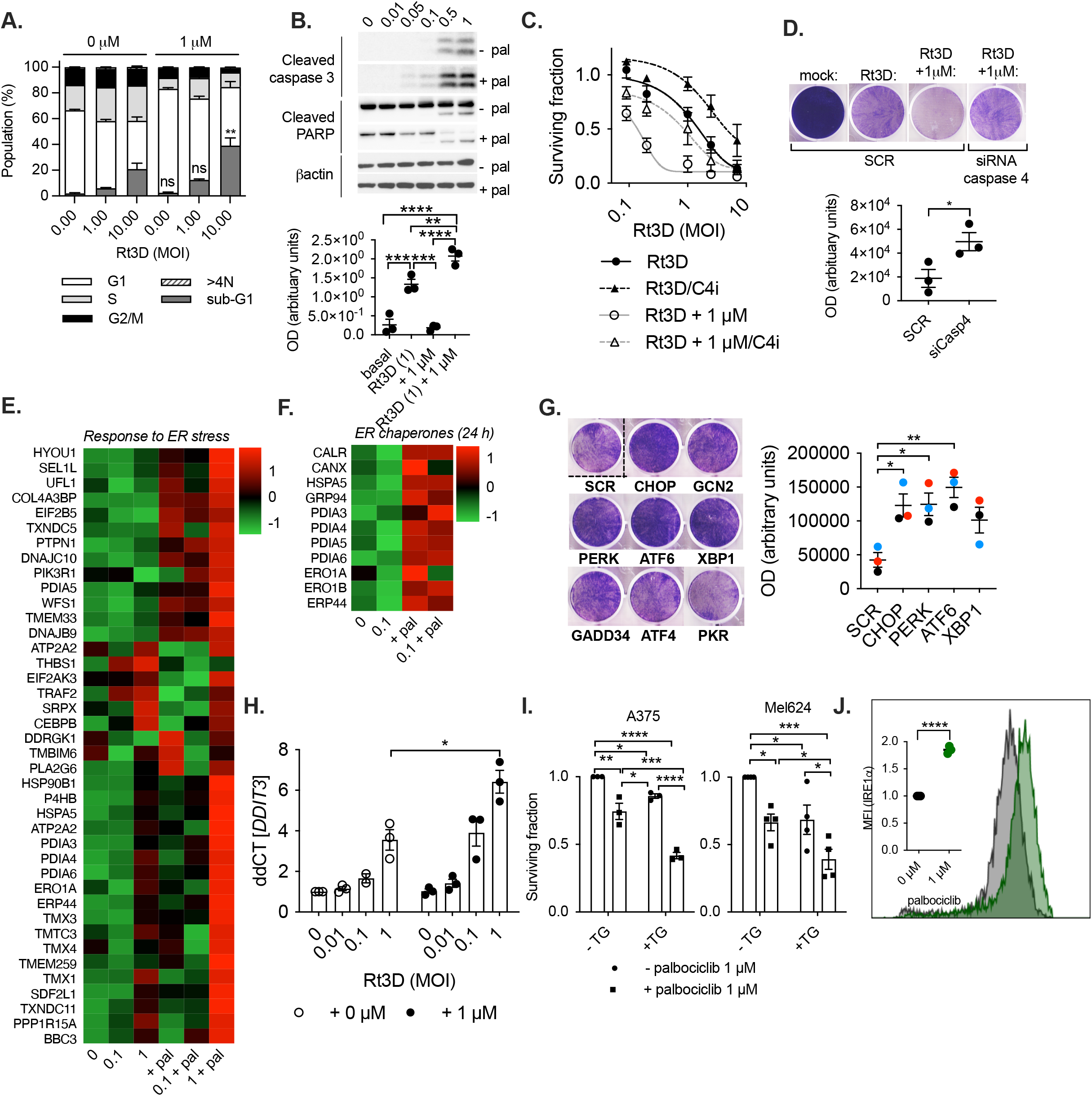
Palbociclib sensitises cells to UPR activation/ER stress and in combination with Rt3D induces an ER stress signature. **A**, FACS analysis of A375 cells treated with Rt3D +/-palbociclib (1 μM) stained for PI at 48 hours (± SEM, *n* = 3). **B**, Western blots showing apoptotic markers in A375 cells treated with Rt3D (0.01-1) +/-palbociclib (1 μM) at 48 hours (representative for 3 biological replicates, quantification is shown below). **C**, Cell viability in A375 cells treated with Rt3D +/-palbociclib (1 μM) +/-caspase-4 inhibitor Z-YVAD-FMK, by MTT (72 hours) (± SEM, *n* = 3). **I**, siRNA silencing of caspase 4 in A375 cells treated with Rt3D (0.1) plus palbociclib (1 μM) by crystal violet (with quantification for 3 biological replicates below). **E**. Proteomic analysis of A375 cells treated with Rt3D (0.1-1) in combination with palbociclib (1 μM) showing up-regulated (red) and down-regulated (green) proteins by combination treatment categorised under the gene ontology (GO) term ‘response to endoplasmic reticulum stress’ (48 hours). **F**. Proteomic analysis of A375 cells treated with Rt3D (0.1) in combination with palbociclib (1 μM) showing ER chaperones (24 hour). **G**, siRNA knockdown of UPR components in A375 cells treated with Rt3D (0.1) plus palbociclib (1 μM) by crystal violet (left) and quantatative analysis for 3 independent experiments (right). **H**, RT-qPCR of CHOP at 48 hours following Rt3D + palbociclib (1 μM) in A375 (± SEM, *n* = 3). **I**, Cell viability in A375 and Mel624 cells treated with thapsigargin (TG, 0.04 μM) +/-palbociclib (1 μM) by MTT assay (± SEM, *n* = 3). **J**, A375 cells containing an IRE1α endonuclease reporter was treated with palbociclib (1 μM) for 72 hours prior to FACS analysis. Inset shows mean fluorescence intesnsity (MFI). (± SEM, *n* = 3).

Caspase-4 is localised to the endoplasmic reticulum (ER) and can be activated in response to ER stress-inducing agents^9^. Therefore, to investigate the possibility that Rt3D-palbociclib therapy induces ER stress, proteome data were analysed under the GO term ‘Response to ER stress’. This showed altered expression of proteins relating to ER stress in the context of Rt3D-palbociclib combination therapy (**Fig. 4E**). Interestingly, proteomic data at an early timepoint (24 hours) also indicated palbociclib therapy-associated elevation of ER chaperones, such as calreticulin, calnexin, GRP78/BiP (HSPA5) and protein disulphide isomerases (**Fig. 4F**).

Rt3D is known to induce ER stress^1,10–12^, linked to remodeling of ER membranes^13,14^, activating unfolded protein response (UPR) signaling via three effector pathways: IRE1α-XBP1, PERK-eIF2α-ATF4 and ATF6^15^. These lead either to restoration of normal protein homeostasis or apoptosis^15^. Therefore, we investigated the importance of these UPR components in the efficacy of combined Rt3D-palbociclib by trying to rescue the cell death phenotype with siRNA. CCAAT-enhancer-binding protein homologous protein (CHOP) induces death due to ER stress^16^. In line with our data on caspase-4, CHOP knockdown also rescued death due to combination treatment, as did PERK and ATF6 knockdown (**Fig. 4G, Fig. S4D**). Rt3D-palbociclib-mediated upregulation of CHOP was confirmed by qPCR (**Fig. 4H**).

Previous studies have also suggested linkages between CDK4/6 inhibition and ER stress^17,18^. Therefore, we next tested if palbociclib could sensitise cells to ER stress-induced cytotoxicity. Palbociclib enhanced cell kill due to the UPR activator, thapsigargin (TG), confirming that it can behave as an ER stress sensitiser (**Fig. 4I**). ER stress has been demonstrated to induce recruitment of XBP1 and transcriptional machinery to the *ifnb1* promoter during PRR activation, resulting in synergistic IFNβ expression in a study using mouse macrophages^19^. Palbociclib increased an IRE1α endonuclease reporter signal, supporting the involvement of the IRE1α-XBP1 branch in the palbociclib-associated ER stress signature (**Fig. 4J, S4E-G**).

### ER stress enhances RNA sensor-driven interferon responses

Due to the ER stress signatures and dsRNA response induced by Rt3D-palbociclib, and the importance of both these pathways in cell death, we suspected that the combination was exploiting cross-talk between UPR signaling and cytoplasmic dsRNA sensing, to mediate increased IFN response and cytotoxicity. Such UPR-RNA cross-talk has been previously described in immune cells^19–21^. We next tested non-viral tool compounds specific to ER stress/UPR activation (thapsigargin) and RNA sensor activation (poly I:C, 3p-hpRNA), to further investigate UPR-RNA cross-talk in tumour cells under alternative treatment conditions. Co-treatment with thapsigargin and poly I:C, either transfected (activating the cytoplasmic dsRNA sensor) or untransfected (acting via endosomal TLR3), enhanced cell kill relative to single agents (**Fig. 5A, B, Fig. S5A**). In response to escalating doses of poly I:C, transcriptional expression of cytoplasmic RNA sensors and type I/III interferons plateaued, suggesting saturation of signaling capacity. However, combining poly I:C with thapsigargin facilitated increased expression of RIG-I (*DDX58*), MDA5 (*IFIH1*), *IFN*α, *IFN*β, *IL-28, IL-29* and CHOP (*DDIT3*), mediating increased downstream signaling from escalating levels of input RNA agonist (**Fig. 5C, Fig. S5B**). The RIG-I agonist, 3p-hpRNA, combined with thapsigargin, induced similar transcriptional changes to those observed for poly I:C (**Fig. 5D, Fig. S5C, D**), as well as increased levels of expression of the ERV *MLT1C49* (**Fig 5E**). Taken together, these data demonstrate generalizable crosstalk between ER stress and RNA agonism in tumour cells resulting in augmented transcription of RNA sensors, amplification of IFN expression and enhanced cell death.

**Figure 5.**
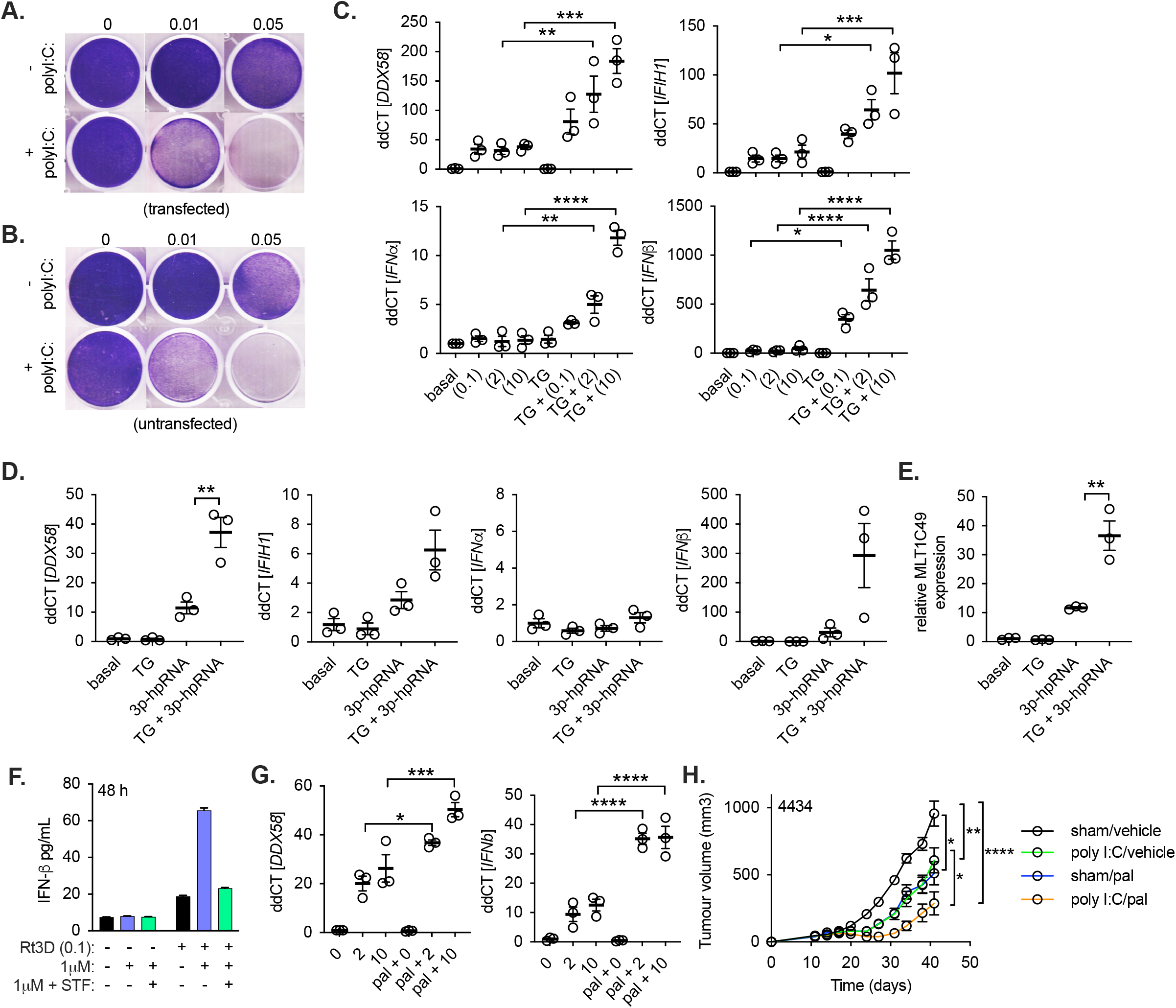
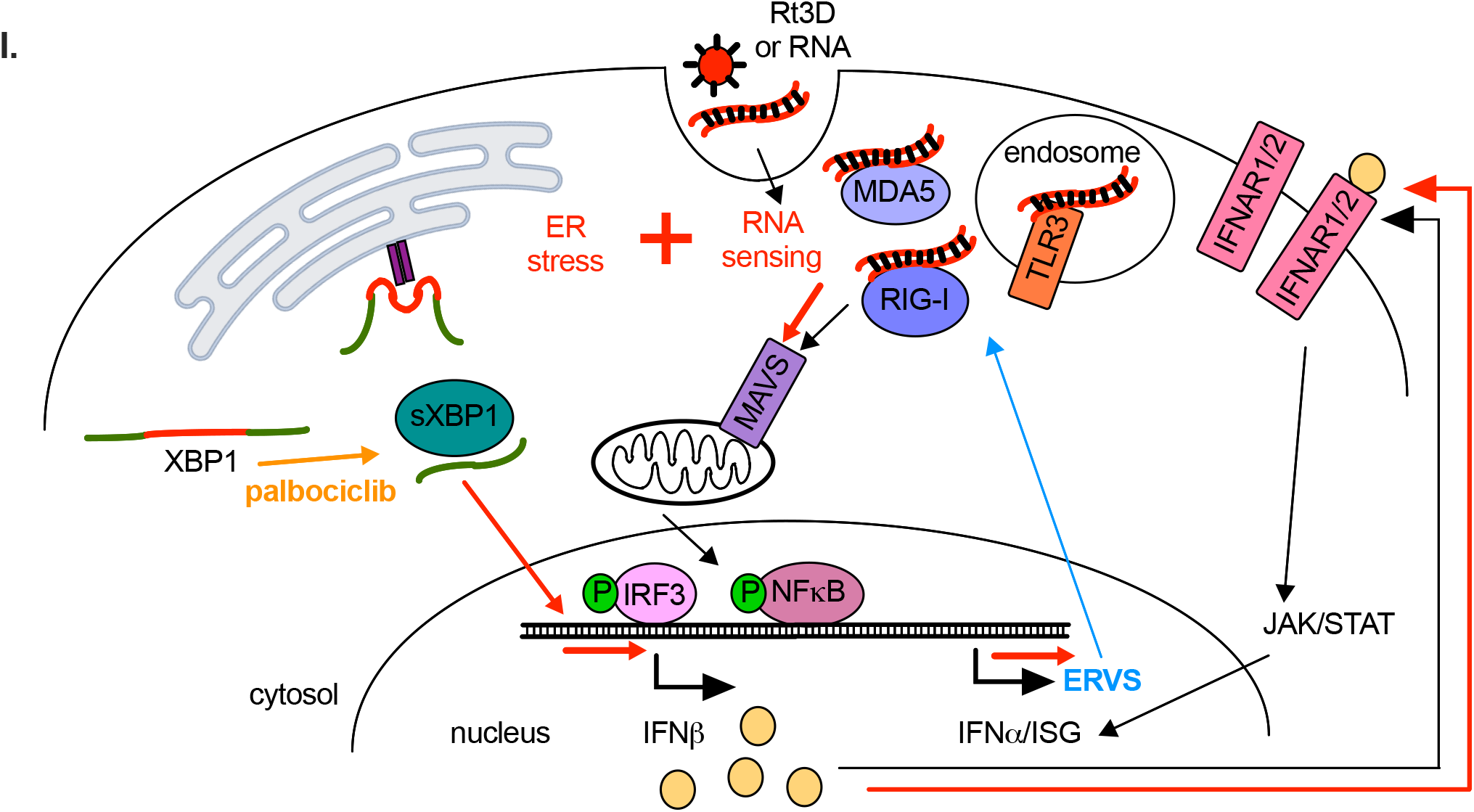
Cross-talk between RNA sensors and ER stress/UPR activation enhances RNA sensing and IFN response. **A**, A375 cells treated with transfected or **B**, untransfected poly I:C (10 μg/mL), combined with thapsigargin (TG, 0.01, 0.05 μM) and cell viability shown by crystal violet assay. **C**, RT-qPCR of RIG-I (*DDX58*), MDA5 (*IFIH1*), *IFN*α *IFN*β in A375 cells treated with transfected poly I:C (0.1, 2, 10 μg/mL), +/-thapsigargin (TG, 0.05 μM) at 48 hours. **D**, RT-qPCR of indicated gene expression in A375 cells treated with RIG-I agonist 3p-hpRNA (1 μg/mL) +/-thapsigargin (0.05 μM) at 48 hours. **E**, RT-qPCR of the ERV *MLT1C49* in A375 cells treated with 3p-hpRNA +/-thapsigargin at 48 hours. **F**. IFNβ measured by ELISA in cell-free supernatant from A375 cells treated with Rt3D (0.1 MOI) -/+ palbociclib (1 μM) +/-IRE1α inhibitor (STF-083010). Data are representative of 2 independent experiments. **G**, RT-qPCR of RIG-I (*DDX58*) and *IFN*β in A375 cells treated with 1 μM palbociclib (+) +/-poly I:C (2, 10 μg/mL) at 48 hours (± SEM, *n* = 3). **H**, Tumour volumes for C57BL/6 mice bearing 4434 tumours treated with a single intratumoural injection of poly I:C (50 μg) in combination with palbociclib (90mg/kg). PCR data are all ± SEM, *n* = 3. + 0 μM

To further test the linkage between UPR signaling and IFN response in Rt3D-palbociclib-treated cells, we evaluated effects of drugs that modulate the UPR on Rt3D-palbociclib-induced IFNβ production^12^. IRE1α inhibition reduced the elevated IFNβ seen with Rt3D-palbociclib therapy (**Fig. 5F, Fig. S5E**). Together with knockdown data (**Fig. 3E, Fig. 4G**), these experiments support ER stress/UPR activation plus RNA sensor agonism through combined Rt3D-palbociclib, as a key driver of interferon signaling and enhanced anti-cancer efficacy in response to dsRNA sensor activation.

Finally, we asked whether palbociclib sensitises cells to RNA in a non-viral context. Indeed, poly I:C/palbociclib significantly amplified RIG-I (*DDX58*) and IFNβ expression above levels seen with palbociclib alone (**Fig. 5G**). Returning to tumour therapy, this combination reduced tumour burden *in vivo*, demonstrating that palbociclib treatment can more widely sensitise tumours to RNA agonism, with therapeutic benefit (**Fig. 5H**). A summary model for the interactions between RNA sensing and ER stress, as addressed in this study, is shown in **Fig. 5I**.

## Discussion

Boosting innate immune effector function by targeting nucleic acid sensors, linked to subsequent priming of antigen-specific T cell-mediated anti-tumour immunity, is currently an exciting area of anti-cancer research, with STING, TLR3 and RIG-I agonists currently under therapeutic investigation^22–24^. Oncolytic viruses can also be classified as nucleic acid sensor agonists, but they have more pleiotropic effects including inducing ER stress and UPR activation. ER stress due to Rt3D is well described^11,25,26^, and we have previously exploited this therapeutically^1,12^. In this study, ER stress/UPR signalling plus RNA agonism potentiated IFN signalling and enhanced cell death and immunogenicity in tumour cells, heralding significant combinatorial opportunities.

We are unaware of any previous description of CDK4/6 inhibition as an ER stress sensitizer. Palbociclib-treated young rats and CDK4-knockout mice display insulin-deficiency, defective pancreatic beta-cell function and increases in ER stress markers^17,18^. Beta-cells are sensitive to perturbation of ER stress, with these studies phenocopying observations in PERK-knockout mice^27^. Consistent with these findings, we confirm that single-agent palbociclib increased ER stress-associated proteins and the IRE1α endonuclease reporter signal in treated tumour cells.

In addition to the ER stress signature seen with Rt3D-palbociclib, we observed a potent increase in the IFN response within tumour cells, both at the transcriptional and protein levels, with an escalation in the expression of RNA sensors. Hence, induction of the UPR goes beyond protein homeostasis, interacting with PRR signaling and enhancing inflammatory responses; this is the first demonstration, to our knowledge, of such cross-talk between these pathways in tumour cells and its immunological consequences. Importantly, this cross-talk extends beyond the use of OV and a targeted small molecule as a therapy combination, since enhanced UPR signalling using thapsigargin also liberated interferon gene expression to poly I:C dsRNA in melanoma cells. Furthermore, IRE1α inhibition was able to reduce Rt3D-palbociclib-induced IFN-beta protein, further demonstrating the link between UPR and IFN responses. Research investigating UPR and PRRs crosstalk has previously been restricted to immune cells; it has not previously been described in the context of cancer therapeutics, which has broad implications for the clinical development of agents including nucleic acid sensor agonists and oncolytic viruses. From data in this study and others, it is apparent that ER stress in and of itself does not induce type-I IFNs^21,28–30^.

Spliced-XBP1 (by the endonuclease IRE1α) has, however, been shown to bind to an enhancer site of the *ifnB1* gene^19^, with numerous studies confirming spliced-XBP1 boosts IFN-beta only in combination with certain PRR activators^19,20,25,30,31^.

As with the tool compound thapsigargin, palbociclib sensitised cancer cells to ER stress which, in combination with PRR activation, also led to increased IFN expression. This combination translated to therapy, as shown by the combination with poly I:C in vivo, illustrating the capacity of palbociclib to amplify the IFN response in a non-viral context.

In this study, we addressed the consequences of the observed increased IFN on tumour cell-intrinsic immunogenicity. Increased HLA expression by Rt3D-palbocilcib was observed both in RNA sequencing and proteomic data and confirmed on the cell surface by FACS analysis. MHC II expression on tumour cells has been previously reported and associated with a therapeutic response to PD-1^32^. Expression of both HLA class I and II was reduced with inhibition of JAK/STAT signaling, demonstrating a link between increased IFN signaling and antigen presentation. CDK4/6i has previously been reported to increase antigen presentation^33^, consistent with the findings in this study.

CDK4/6i has been shown to induce dsRNA-IFN responses through downregulation of the E2F target DNMT1 and expression of endogenous retroviral (ERV) elements^8,33^. Indeed, we observed marked modification to DNA including down-regulation of DNMT as a result of palbociclib treatment, although no changes were seen in DNA methylation status (data not shown). We did, however, observe a marked downregulation in histones. Analysis of chromatin accessibility, by methods such as ATAC sequencing may provide more insight into the palbociclib-driven effects on histone loss, and the consequences on the transcriptome (including ERV expression). By screening a selection of ERVs, we observed enhanced STAT1-driven transcription of select ERVs. This was not specifically a virus-mediated effect, because ERV expression was also accomplished by the treatment of RIG-I stimulation and amplified in combination with ER stress (thapsigargin).

Upregulation of shared tumour-associated antigens such as MAGEA4, as well as enhanced STAT1-driven ERV transcription are likely to be beneficial in promoting anti-tumor immune responses^34–36^. Herein, we demonstrate that peptides originating from CTA and ERV elements can be presented by HLA class I molecules on tumour cells. Whether these peptides and their expression reflects alterations in functionally relevant tumour-associated antigens, which contribute to adaptive immune therapy, is currently not known. Despite the presence of ERV peptides on the immunopeptidome, no changes were seen in the RNA expression of these ERVs with Rt3D-palbociclib treatment. This phenomenon was also observed for MAGEA4 expression, and suggests that post-transcriptional processing events (such as proteasome/immunoproteasome switching) may be responsible for the appearance of these peptides associated with HLA I. Overall, the wider functional immune consequences of the tumour cell-intrinsic immune activation by CDK4/6 inhibition and double-stranded RNA sensor agonism described in this study, requires further investigation. Key questions, including i) the extent to which the immune system plays a role in Rt3D-palbociclib combination therapy in vivo, ii) the effects of Rt3D-palbociclib on the tumour immune microenvironment, and iii) whether the potentially antigenic peptides we have found presented by Rt3D-palbociclib-treated cells (eg derived from ERV/CTA) are important for in vivo therapy, are currently being addressed in our laboratory.

This study demonstrates that cross-talk between RNA sensing and ER stress pathways can increase tumour cell death, particularly in the application of the clinical therapies CDK4/6 inhibition and RNA agonism/oncolytic virotherapy. These interventions also increase the immunogenicity of tumour cell death, both in terms of innate interferon production, and adaptive presentation by HLA class I peptides of potential TAA. Further work is currently underway to fully define the functional immune consequences of this novel combinatorial approach. Our data supports clinical translational development of combined RNA sensing and ER stress pathway engagement. In this regard, both palbociclib and Rt3D are excellent candidates, given the fact that the former is an approved anti-cancer agent and the latter has been the subject of multiple clinical trials demonstrating safety and tolerability.

## Materials and Methods

### Cell lines and therapeutic agents

Neat stocks of 1 × 10^9^ TCID_50_ Rt3D were diluted 1:10 in DMEM for single use aliquots and stored at -80ºC. Palbociclib was kindly provided by Pfizer.

A375, Mel624, MeWo, A2058, Fadu, HN5 and Cal27 were obtained from stocks within Kevin Harrington’s team at the ICR. D04 were obtained by generous donation from R. Marais, CRUK Manchester Institute. T47D, T47D-pR, MCF7, MCF-7-pR were obtained by generous donation from N. Turner, ICR London and cultured in RPMI (no phenol red) plus oestradiol (1 nM) with the addition of 0.5 µM palbociclib for resistant (pR) cell lines. All other cell lines were obtained from stocks within Kevin Harrington’s team at the ICR. DMEM or RPMI was supplemented with 5% or 10% (v/v) FCS, 1% (v/v) glutamine, and 0.5% (v/v) penicillin/streptomycin (ICR, CSSD, UK).

### Fucci and reporter cells

Cells were infected with Fluorescent Ubiquitination-based Cell-Cycle Indicator (FUCCI) cell-cycle vectors. G1 red expresses mCherry hCdt1(30-120aa); S– G2–M cyan expresses AmCyan hGeminin(1-110aa). Sequences derived from pRetroX-G1-Red and pRetroX-SG2M (Clontech) were cloned into pHR lentiviral vectors under puromycin or blasticidin selection. Lentiviral packaging was performed in 293T cells using MD2.g and psPAX2. A375 cells containing an IRE1α endonuclease reporter cells were generated as previously described (McLaughlin et al).

### Library screen and cell titre glo assays

Cells were plated at 500 cells per well in 20 µL, in 384 well plates using a Thermo Scientific multidrop combi and incubated overnight. Readily available small molecule screen plates 1 (11) and 2 (12) were used compromising a range of different concentrations of 80 drugs (Prof. Chris Lord, ICR). Medium containing 20 µL of the drug ranging from 1-1000 nM was added to the cells using a Hamilton microlab star. Cells were infected with Rt3D in 5 µL per well, 2 hours after the addition of the compounds. 72 hours after treatment, cell viability was measured by CellTiter-Glo Luminescent Cell Viability Assay (CTG, Promega, UK). 25 µL of CTG was added per well and incubated at 37ºC for 10 minutes. Luminescence was measured on a Wallac Victor 2V plate reader.

### Screen analysis

In each plate, 8 dose points were used per compound. Combined with 4 doses of Rt3D, this was a comprehensive 8 × 5 matrix screen. Rt3D cytotoxicity levels between plates were compared for consistency. The effect of the drug-virotherapy combination was ranked as ‘therapy effect’ Z-scores. In brief, Z scores are a normalized value between untreated (library alone) versus treated (virus plus library) for each dose used. A threshold value <-2 was used to define highly significant sensitization (values of >2 would indicate combination therapy resistance). The data have been normalized to untreated or virus alone in each case.

### MTT assays

Cells were seeded between 5 × 10^3^ – 1 × 10^4^ cells per well in 96-well plates overnight prior to treatment with palbociclib at indicated doses (Pfizer), thapsigargin (T9033, Sigma), 5-Azacytidine (A2385, Sigma), poly I:C (P1530, Sigma) and/or infected with Rt3D at indicated MOI. At the desired time point, 20 µL of 5 mg/mL 3-(4,5-dimethylthiazol-2-yl)-2,5-diphenyltetrazolium bromide (MTT) was added per well and incubated for 2-4 hours. The supernatant was removed, and the formazan was solubilised in DMSO. Absorbance was read in a plate reader at 550 nm.

### Crystal violet and SRB assays

Cells were seeded in 6-well plates at 3-6 × 10^5^ cells per well overnight and treated as described. For crystal violet assays, medium was removed, and cells were washed in PBS. 2 mL of crystal violet was added per well and incubated for 20 minutes, after which time it was removed, and the plates washed. For SRB assays, cells were fixed by adding an equal volume of 10% TCA for 1 hour. Plates were then washed in tap water and left to dry prior to staining with 0.057% SRB (wt/vol) in 1% acetic acid overnight. Plates were then washed in 1% acetic acid and left to dry.

### Clonogenic assays

Cells were seeded in 6 well plates at 7.5 × 10^3^ cells per well overnight. After treatment with palbociclib or Rt3D, cells were incubated for 2 weeks with renewal of cell medium every 4 days (replacing drug but not virus). Colonies were measured by crystal violet assay.

### Cell cycle assays

Cells were seeded in 6-well plates at 3 × 10^5^ cells per well overnight prior to treatments. Post-treatment, cells were harvested, washed and then fixed with ethanol overnight. The following day, cells were pelleted out of ethanol, and stained with propidium iodide (PI) (40µg/mL) in the presence of 10 μg/mL RNase A (R4875, Sigma) prior to analysis on the BD LSR II analyser.

### FACS analysis

Cells were seeded in 6-well plates at 3 × 10^5^ cells per well overnight prior to treatments. At the desired timepoint, cells were harvested and spun at 1200 rpm, 5 mins. Cells were resuspended in antibodies anti-human HLA-A,B,C (311403, biolegend), HLA-DR,DP,DQ (361709, biolegend) with viability dye (65-0865-14, eBioscience) for 30 mins. Cells were washed in PBS + 5% FBS and fixed in FluoroFix buffer (422101, BioLegend) prior to analysis on the BD LSR II analyser.

### In vivo experiments

CD1 nude mice or C57BL/6 mice (Charles Rivers, Kent, UK) were subcutaneously injected with 3 × 10^6^ A375 cells or 4 × 10^6^ 4434 cells respectively, suspended in PBS in the right flank. Once tumors were established to ∼6 mm in diameter, mice were allocated treatment groups stratified by tumor size. Mice bearing tumors were treated daily with 60-90 mg/kg palbociclib (Pfizer) or vehicle (0.05 N sodium lactate buffer, pH 4.0) by oral gavage. Drug administration continued daily for 2 weeks. A Rt3D injection of 1 × 10^6^ pfu dissolved in PBS (or a PBS sham) was administered as an intra-tumoral injection 3 days after drug administration commenced. Poly I:C was given as a single intratumoural injection of 50 μg (tlrl-pic, Invivogen). Established tumor volumes were measured at least twice weekly using Vernier calipers and the tumor volume was estimated from the formula: V = 0.5 × (length × width^2^). All experiments were carried out in compliance with the NCRI guidelines, with animals judged to have failed treatment if tumor diameter exceeded 10mm (A375), or 15mm (4434).

### Immune profiling of tumours

C57BL/6 mice (Charles Rivers, Kent, UK) were subcutaneously implanted with 3 × 106 MOC1 murine cells suspended in 0.1 mL PBS per flank. Tumours were allowed to grow to 6-8 mm before mice were allocated treatment groups stratified by tumour size. Mice bearing tumours were treated daily with 100 mg/kg palbociclib (kindly provided by Pfizer) or vehicle (0.05 N sodium lactate buffer, pH 4.0, refer to palbociclib stocks) by oral gavage. A Rt3D injection of 5 × 10^6^ pfu dissolved in PBS (or a PBS sham) was administered as an intra-tumoural injection 3 days after drug administration commenced. Tumours were harvested (5 mice per group) and analysed for tumour-infiltrating lymphocytes by FACS. The timepoints of collection were day 3, 7 and 10 after Rt3D injection. Tumours were harvested and minced with scissors in digestion mix (0.01% trypsin, 2.5 mg/mL collagenase, 2 mg/mL dispase and 1 mg/mL DNAse in RPMI) and incubated at 37°C for 30 minutes. Thereafter, samples were kept on ice. Suspensions were passed through a 70 ^μ^m strainer using a 2.5 mL syringe plunger and washed through with RPMI + 5 mM EDTA until only connective tissue remained. Samples were centrifuged at 1500 rpm, for 5 mins at 4°C, and transferred into a V-well 96-well plate. Samples were stained in FACS buffer (PBS + 5% FCS) for 30 mins on ice and protected from light, with the following extracellular antibodies; CD3 (100218), CD4 (100406), CD8 (562315), CD45 (103125), Ly6G (127633), CD11b (101205), Ly6C (128015), CD11c (117327), MHC II (107621), PD-L1 (558091) F4/80 (123113) from BioLegend and viability dye (65-0865-14) from Thermo Fisher Scientific. Cells were then washed in FACS buffer and permeabilized and stained with intracellular antibody to FOXP3 (48-5773-80) or Ki-67 (69-5698-80) both from Thermo Fisher Scientific. Samples were then washed and fixed (1-2% PFA) prior to analysis of tumour-infiltrating lymphocytes by flow cytometry. Tumours were weighed on collection and counting beads were added when running the analysis to calculate cells per mg of tumour.

### Western blotting

Samples harvested for western blotting were stained with phospho^(ser780)^-Rb, phospho^(ser807/811)^-Rb, CDK4, Cyclin D1, Cyclin E1, CyclinA2, phospho-EIF2_α_ (#3398), caspase-3 (#9665), HMGB1 (#3935), phospho-STAT-1 (#9167), RIG-I (#3743) (New England Biolabs, UK), PARP-1 (F-2) (sc-8007, Santa Cruz Biotechnology), anti-DNMT (ab92314, abcam), MDA5 (AT113, ALX-210-935, Enzo), or anti-beta actin [AC-15] (ab6276, abcam, UK). µ1C 10F6 (reovirus) was deposited to the DSHB by Dermody, T.S. (DSHB Hybridoma Product 10F6 (reovirus)).

### ZVAD experiments

Cells were seeded in 96 well plates at 5 × 10^3^ cells per well overnight. A inhibitor sampler pack which contains caspase inhibitors of 1, 2, 3, 4, 6, 8, 9, 10, 13 and a pan-caspase inhibitor (R&D systems), was used on cells treated with combination therapy to observe rescue from cells treated without inhibitors. The caspase 4 inhibitor (Z-YVAD-FMK, FMK005, R&D systems) was used in further experiments at the dose of 50 µM. Cell survival was measured by MTT assay 72 hours later.

### siRNA experiments

Cells were seeded in 12 or 6-well plates at 5 × 10^5^ or 3 × 10^5^ cells per well respectively overnight. Cells were transfected with siRNA (Qiagen) using Lipofectamine RNAiMAX transfection reagent (Thermo Fisher) as per manufacturers instructions. 24 hours later media was replaced containing therapy and cell survival was measured 48-96 hours later by crystal violet assay.

### Viral replication assays

Cells were plated at 1 × 10^5^ in 24-well plates and treated the following day with palbociclib and/or Rt3D at a dose of MOI 5 (to ensure infection of all cells). After another 2 hours, cells were washed twice with media and the inhibitors replaced. At 4, 24, and 48 hours after infection, cells were harvested into the media and stored at −80°C prior to processing. The lysate was subjected to 3× freeze/thawing between −80 and 37°C, centrifuged at 13,000 rpm for 5 minutes and the supernatant was used to titre for Rt3D by TCID50 assay on L929 cells.

### RT-qPCR

RNA was extracted from samples using RNeasy kit (Qiagen, UK), and cDNA was synthesised using SensiFAST cDNA synthesis kit before amplification against transcripts by qRT-PCR kit with SYBR green (Bioline). Alternatively, transcripts were amplified directly on RNA samples using a one-step qRT-PCR kit with SYBR green (Bioline). Primers used against *DDIT3* (CHOP) were commercially available QuantiTect primer assays (Qiagen, UK). Please refer to **table S2** and **S3** in the supplementary material for the sequences of other primers used. Relative gene expression was calculated with the 2-ddCT method using 18S or beta actin as a house-keeping gene.

### ATP release

Cells were plated at 3 × 10^5^ cells per well in 6-well plates and incubated overnight prior to treatment the following day. Cell supernatants were collected at the desired timepoints and centrifuged at 2000 rpm for 4 mins. 200 μL of cell-free supernatants were mixed with 50μL CellTiter-Glo Luminescent Cell Viability Assay (Promega, G7571) in white 96-well plates (Greiner, 655075). After 10 minutes incubation at room temperature, luminescence was measured on a Victor 2V plate reader.

### Confocal microscopy

Cells were seeded in glass-bottom dishes at 3 × 10^5^ cells per well overnight prior to treatments. After treatment cell medium was removed, and cells were fixed in 10% formalin for 5-10 minutes. Samples were then washed and then blocked (with PBS + 1 % BSA, 2 % FBS and 0.05% Sodium Azide). Samples were stained with calreticulin primary antibody (PA3-900, Thermofisher, UK), secondary antibody conjugated to FITC (A-11070 anti-rabbit Alexafluor 488, Invitrogen, UK), Hoechst 33342, and imaged by confocal microscopy by Z stacking. 3 fields of view were taken for 3 independent repeats and CRT expression was quantified relative to cell number using cell profiler.

### Phagocytosis assays

The methodology for this assay has been previously described^6^. PBMCs were harvested from K3EDTA anti-coagulated whole blood by density gradient centrifugation with Lymphoprep and plated in 12 well plates at 2-4 × 10^6^ cells per well in 10% RPMI + 10 ng/mL human CSF (300-25, Peprotech, UK). Medium was replaced every 2-3 days until adherent populations of macrophages had established, and these were used in experiments 8-10 days later. Tumour cells were stained with 20 ng/mL of pHrodo-SE (P36600, Thermofisher) for 30 minutes then washed and co-cultured with the macrophages. 1-2 hours after co-culture, samples were washed and stained for CD11b conjugated to FITC (#11-0118-42, Thermofisher), then fixed and analysed on the BD LSR II. Separately stained macrophages or treated cells labelled with pHrodo-SE served to determine cut-offs for phagocytosis. In 1 × 10^4^ collected events, phagocytosis percentage was determined by the double-stained CD11b+ pHrodo-SE+ cells from the CD11b+ population above the threshold set in the controls.

### Whole proteome LC-MS/MS Analysis

Cells were seeded in 6 cm dishes at 9 × 10^5^ cells per well overnight prior to treatments and collected at indicated time-points by trypsinising cells and washing 3 times in PBS. Pellets were stored at −80°C prior to processing. Cell pellets were lysed in 5% SDS / 100 mM TEAB (tetraethylammonium bromide, Sigma) by probe sonication and heating at 90LC for 10 min. Proteins were reduced by TCEP (Tris(2-carboxyethyl) phosphine, Sigma), alkylated by iodoacetamide (Sigma), and purified by trichloroacetic acid (Sigma) precipitation. The protein pellet was resuspended in 100 mM TEAB buffer and digested by trypsin (Thermo). 40 µg of protein digest was labelled by TMT11plex (Thermo Fisher), and 11 samples were pooled, dried in SpeedVac. The mixture was fractionated on an XBridge BEH C18 column (2.1 mm i.d. x 150 mm, Waters). Fractions were collected at every 30 sec and then concatenated to 28 fractions and dried in SpeedVac. 1/3 of peptides were injected for on-line LC-MS/MS analysis on the Orbitraip Fusion Lumos hybrid mass spectrometer coupled with an Ultimate 3000 RSLCnano UPLC system. Peptides were first loaded on a PepMap C18 nano trap (100 µm i.d. x 20 mm, 100 Å, 5µ), and then separated on a PepMap C18 column (75 µm i.d. x 500 mm, 2 µm) over a linear gradient of 6.4 - 30.5% CH_3_CN/0.1% formic acid in 90 min / cycle time at 120 min at a flow rate at 300 nL/min. All instrument and columns were from Thermo Fisher. The MS acquisition used MS3 level quantification with Synchronous Precursor Selection (SPS5) with the Top Speed 3s cycle time. Briefly, the full MS survey scan in Orbitrap was m/z 375 – 1500. Multiply charged ions (2 – 5) with intensity threshold above 7 × 10^3^ were fragmented in ion trap at 35% collision energy with isolation width at 0.7 Da in quadruple. The dynamic range was 40 s at ±10 ppm. The top 5 MS2 fragment ions were SPS selected with the isolation width at 0.7 Da and fragmented in HCD at 65% NCE and detected in Orbitrap.

### Whole proteome data analysis

Raw spectra were processed using ProteomeDiscoverer v2.4(Thermo Scientific) and searched against FASTA sequence databases containing GENCODE v32^37^ protein sequences, UniProt (2019_05) Reovirus Proteins, translated gEVE database^38^ sequences and cRap contaminates using both Mascot server v2.4 (Matrix Science) and SequestHT with target-decoy scoring evaluated using Percolator^39^. The precursor tolerance was set at 20 ppm, fragment tolerance set at 0.5 Da and spectra were matched with fully tryptic peptides with a maximum of 2 missed cleavages. Fixed modifications included:carbamidomethyl [C] and TMT6plex [N-Term]. Variable modifications included: TMT6plex [K], oxidation [M], and deamidation [NQ]. Peptide results were initially filtered to a 1% FDR(0.01 q-value). The reporter ion quantifier node included a TMT-11-plex quantification method with an integration window tolerance of 15 ppm and integration method based on the most confident centroid peak at MS3 level. Protein quantification was performed using unique peptides only, with protein groups considered for peptide uniqueness. Log2 fold change ratios were calculated for each sample vs time point Basal sample using normalised protein abundances. Ratios and abundance were loaded into Perseus^40^ or further downstream analysis and plotting. Z-score scaling was used to generate heatmaps, proteins were clustered using k-means method and GO enrichment was performed using fisher exact test. Results and RAW spectral files have been uploaded to PRIDE repository^41^ under project accession PXD036540.

### HLA-I peptide capture and TMT labelling

To capture HLA peptides, HLA class I molecules were purified from 100 × 10^6^ cells by immunoaffinity precipitation (IP). Snap frozen cell pellets were lysed in 0.5% CHAPS (Abcam) supplemented with protease and phosphatase inhibitors (Thermo Scientific) for 1h at 4°C. The lysate was centrifuged (30 min., 10000 x g at 4°C) and supernatants used for HLA-I IP in which purified anti-human HLA-A,B,C antibody (W6/32, BioLegend) was crosslinked to Protein G Dynabeads (Thermo Scientific) with Dimethyl pimelimidate dihydrochloride (DMP, Sigma Aldrich). Beads were washed with phosphate buffered saline (PBS) 0.01% Tween 20 (Merck Millipore), incubated with antibody and PBS 0.1% Tween (15 min. at RT), washed twice with PBS 0.01% Tween and twice with 0.2 M Triethanolamine (pH 8.2), prior to crosslinking. Beads and antibodies were crosslinked using 20 mM DMP in Triethanolamine (pH 8.2) (30 min. at RT), and quenched with 50 mM Tris-HCl (15 min. at RT). Beads were washed with PBS 0.1% Tween before incubation with cell lysate supernatants (3h, at 4°C). Beads were washed eight times with cold PBS followed by two washes with cold water prior to peptide elution, in 1ml 0.2 % trifluoroacetic acid on ice. Four rounds of elution were performed on each sample. The beads were reused for a second round of IP using the lysate flowthrough from the first IP. Eluted peptides were pooled, dried, reduced with 5mM TCEP (15 min. at RT) and alkylated with 10 mM iodoacetamide (30 min. at RT). Purified HLA class 1 peptides were labelled by 1mg TMTpro, followed by desalting and fractionation on the high pH spin column (Thermo) as the step elution method by these solvent composition: 10%, 12.5%, 15%, 17.5%, 20%, 25% and 30%, then concatenated to 3 fractions. Samples were dried down by SpeedVac, and resuspended in 0.5%FA before LC-MS analysis.

### Immunopeptidome MS analysis

Samples were analysed by LC-MS/MS on the Orbitrap Fusion Lumos mass spectrometer coupled to a U3000 RSLCnano UHPLC system. All instruments and columns used were from Thermo Fisher. The peptides were first loaded to a PepMap C18 trap (100 µm i.d. x 20 mm, 100 Å, 5 µm) at 10 µl/min with 0.1% FA/H2O, and then separated on a PepMap C18 column (75 µm i.d. x 500 mm, 100 Å, 2 µm) at 300 nl/min and a linear gradient of 4-32% ACN/0.1% FA in 60 min with the total cycle at 90 min. The Orbitrap full MS survey scan was m/z 375 – 2000 with the resolution 120,000 at m/z 200, with AGC (Automatic Gain Control) set at 40,000 and maximum injection time at 50 ms. Ions with charge at 1–4 with intensity above 50,000 counts were fragmented in HCD (higher collision dissociation) cell at 35 % collision energy, and the isolation window at 1 Da. The fragment ions were detected in Orbitrap with AGC at 50,000 and 86 ms maximum injection time. The dynamic exclusion time was set at 30 s with ±10 ppm. The vials were then resuspended in 20% ACN/0.5% FA, diluted to 5% ACN, and followed by another LC-MS analysis with a shorter gradient.

### Immunopeptidome data analysis

Raw MS spectra were converted to mzML format using ThermoRawFileParser^42^. Spectra were searched in a custom NextFlow and OpenMS (v2.6)^43^ pipeline using Comet (2019.01 rev. 5)^44^ and MSGFplus (v2020.07.02)^45^. Results from both search engines were processed by Percolator and merged using the OpenMS tool ConsensusID. The searches were performed against GENCODE v38 Protein Sequences, UniProt (June 2021) ReoVirus Proteins, cRap contaminates, and ERV proteins translated from gEVE database^38^ and hERVd database^46^. Search parameters included minimum peptide length of 8 amino acids and maximum of 15 amino acids with unspecified peptide cleavage. A maximum peptide charge of 7 was allowed. Precursor tolerance was set to 20 ppm. MSGFplus instrument type was set to ‘high_res’ and Comet fragment tolerance was set to 0.01 Da with a bin offset of 0, mass range of 600Da - 2500Da and maximum fragment charge of 5. Fixed modifications included TMTpro (N-term), TMTpro (K), and Carbamidomethyl (C). Variable modifications included Oxidation (M). Results were filtered at a 5% Peptide FDR, and A375 HLA binding was predicted using NetMHC (4.1)^47^. Results and RAW spectral files have been uploaded to PRIDE repository^41^ under project accession PXD036559.

### RNA sequencing

Cells were seeded in 6-well plates at 3 × 10^5^ overnight prior to treatments. 48 hours after treatment cells were harvested using the RNeasy kit (74104, Qiagen) with on-column DNase digestion (79254, Qiagen). Samples were sequenced in-house (at the ICR, London). The STAR alignment software (v.2.5.1b) was used to align reads to the soft masked human reference genome containing all the repeat and low complexity sequences (GRCH38_sm). Custom generated GTF files for TE annotations were downloaded from the TEToolkit website (http://labshare.cshl.edu/shares/mhammelllab/www-data/TEtranscripts/TE_GTF/). Downstream ERV expression analysis was carried out using TETranscript from TEToolkit suite (v2.03), or the SalmonTE (v0.2) pipeline.

### ELISA

A375 cells were seeded at 2 × 10^5^ / mL and treated with 1µM palbociclib then infected with Rt3D for 48 hrs. The production of human IFN_α_ (Mabtech), IFN_β_ (PBL Interferon Source), IL-28 and IL-29, (R & D Systems) in cell free supernatant was determined using matched-paired antibodies according to the manufacturers’ instructions. Optical density absorbance readings were determined at 405 nm absorbance. For UPR inhibitor experiments the compounds STF-083010 (IRE1α inhibitor) was used at 20 µM and IFNβ was measured using the DuoSet ELISA (DY814-05, R & D Systems) as per manufacturers instructions.

### Statistical analysis

T-tests or ANOVA tests were used to make comparisons between groups. For *in vivo* analyses, the area under curve (AUC) was calculated for individual mice. A normality test was performed to verify the data follow a Gaussian distribution. If they do, a T-test or ANOVA test was used followed by a multiple comparisons test, if not, a non-parametric Mann-Whitney test was performed followed by a Dunn’s multiple comparison test. P values were derived where p > 0.05 ns, *p ≤ 0.05, **p ≤ 0.01, ***p ≤0.001, ****p ≤0.0001.

## Acknowledgements

We thank Oncolytics Biotech for providing reovirus. Victoria Roulstone is supported by the Mark Donegan foundation. Martin McLaughlin received funding from the Oracle Cancer Trust. Kevin Harrington is supported by the RM/ICR NIHR Biomedical research centre. We thank Harriet Whittock for help with *in vivo* work produced for this study.

## Figure Legends

### ER stress and double-stranded RNA sensor agonism exert enhanced IFN signalling and expression of JAK/STAT-dependent ERV expression

RNA sensing through Rt3D infection or RNA transfection induces IFNβ expression, which is amplificed by IRE1α mediated XBP1 splicing, achieved by ER stress or palbociclib. JAK/STAT signalling results in expression of ERVs, that may be detected by RNA sensors in a feedback loop.

