## Supplemental figure 1 for "CDK4/6 inhibition and dsRNA sensor agonism co-operate to enhance anti-cancer effects through ER stress and immune modulation of tumour cells"

### Supplementry figure 1.

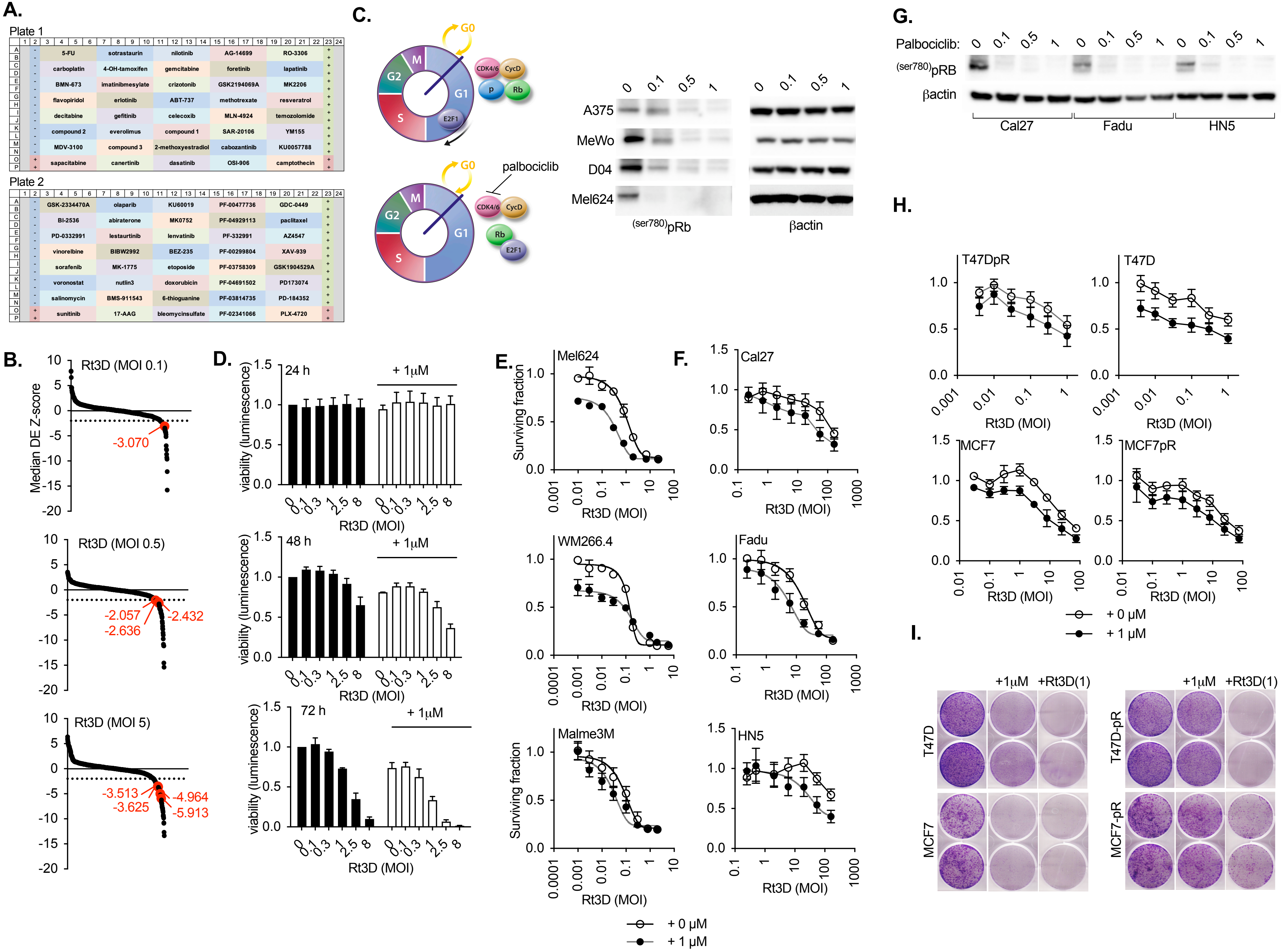

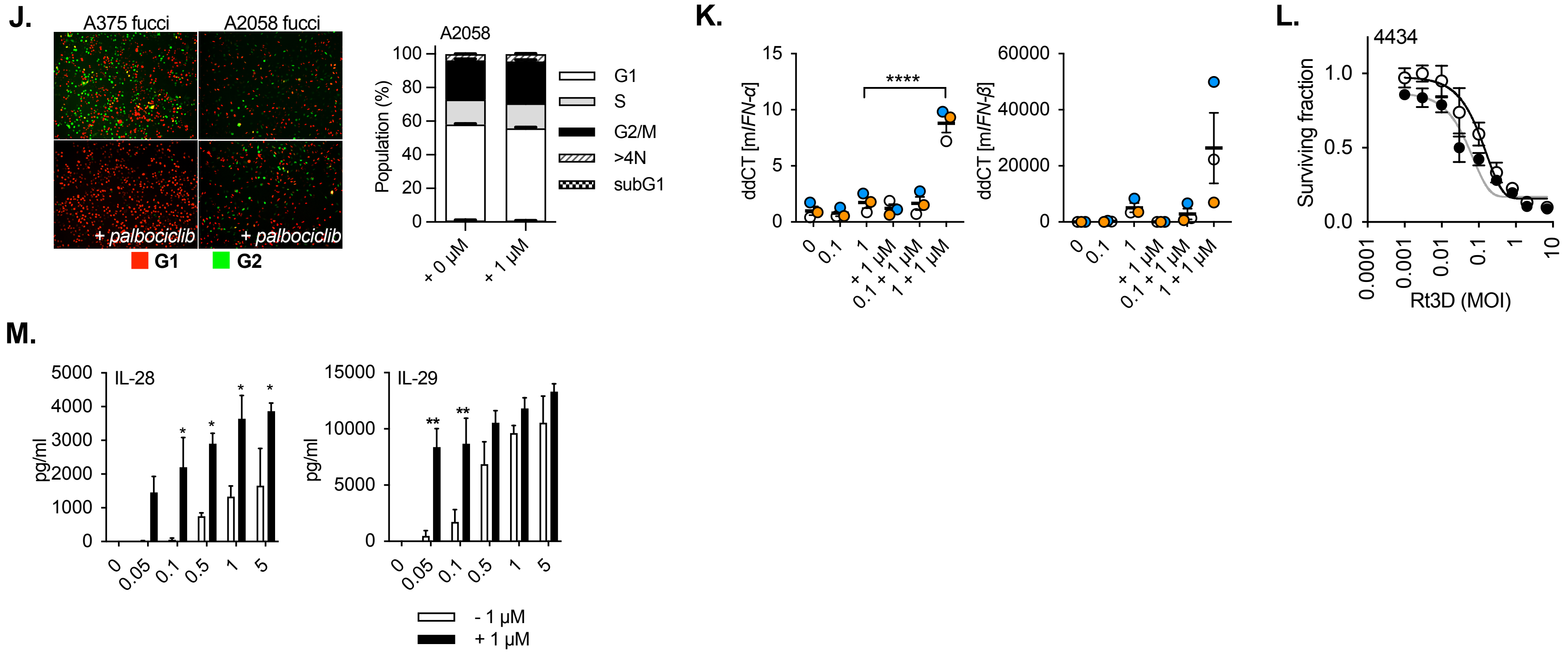


**Supplementary figure 1. A,** Drugs used in the screen are shown in 384-well plate layouts. Controls include medium alone (-), staurosporine (+) or virus alone. Columns 1 and 24 were left blank. **B,** Waterfall plots for A375 BRAF$^{V600E}$-mutant melanoma cells, treated with 80 compounds (at various doses) prior to infection with Rt3D at doses of MOI 0.1-5. Low Z scores (below -2, dashed line) show cell kill sensitisers (palbociclib is shown in red, with relative Z scores). **C.** Palbociclib is a CDK4/6 inhibitor, which maintains Rb hypo-phosphorylation, which renders the transcription factor E2F1 unable to express genes responsible to progress through the cell cycle. The mechanism of action is shown (left). Western blot of pRb treated with indicated doses of palbociclib in a panel of melanoma cell lines at 24 hours (right). **D,** Viability of A375 treated with indicated doses of Rt3D plus palbociclib by cell titre glo assay at 24, 48 and 72 hours. **E,** Cell viability was measured at 72 hours by MTT assay in melanoma cells treated -/+ palbociclib (1 µM) (± SEM, *n* = 3). **f,** Cell viability was measured at 72 hours by MTT assay in head and neck cancer cells treated -/+ palbociclib (1 µM) (± SEM, *n* = 3). **G,** Western blot of pRb treated with indicated doses of palbociclib in a panel of head and neck cell lines at 24 hours. **H,** Breast cancer cells T47D, MCF7 and their palbociclib-resistant counterparts were treated with Rt3D -/+ palbociclib (1 µM) and cell viability was measured at 72 hours by MTT assay (± SEM, *n* = 3). **I,** Colonogenic assays for breast cancer cells that are sensitive (T47D, MCF7) or resistant (T47DpR, MCF7pR) to palbociclib, treated with palbociclib (1 µM), or Rt3D (MOI 1). **J,** Pictomicrographs of fucci versions of Rb wild-type A375 or Rb null A2058 treated with palbociclib (1 µM) at 24 hours (left) and cell cycle analysis in PI stained A2058 cells by FACS (right) (± SEM, *n* = 3). **K,** RT-qPCR of mouse *IFN* and *IFN* in murine BRAF mutant melanoma 4434 cells treated with Rt3D +/- palbociclib (1 µM) at 48 hours (± SEM, *n* = 3). **L,** *in vitro* data display cell viability measured at 72 hours by MTT assay in 4434 cells treated -/+ palbociclib (1 µM) (right) (± SEM, *n* = 3). **M,** IL-28 and IL-29, measured in cell-free supernatant from Rt3D-treated samples (MOI 0.05-0.5) +/- palbociclib (1 µM) by ELISA.
