## Supplemental figure 2 for "CDK4/6 inhibition and dsRNA sensor agonism co-operate to enhance anti-cancer effects through ER stress and immune modulation of tumour cells"

**Supplementry figure 2.**

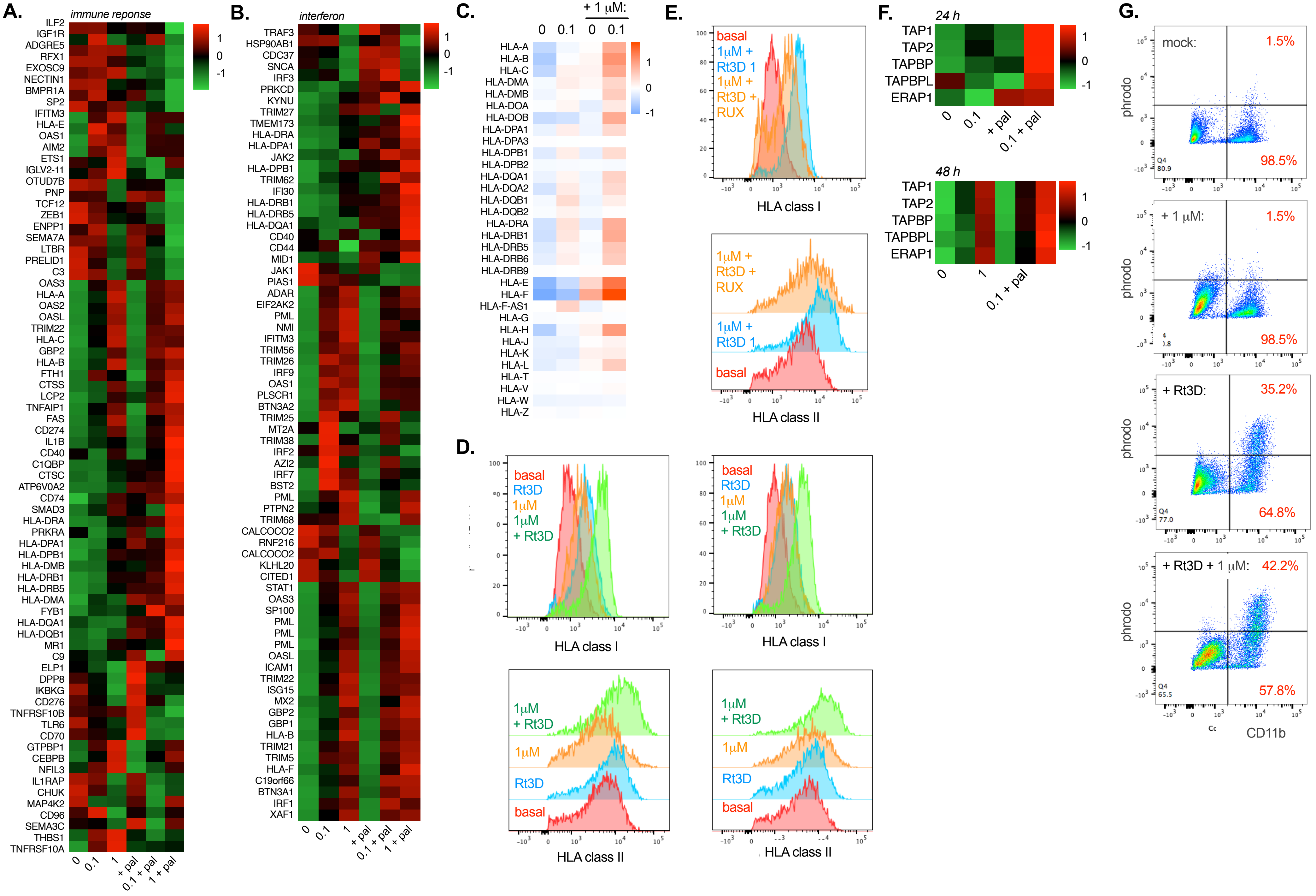

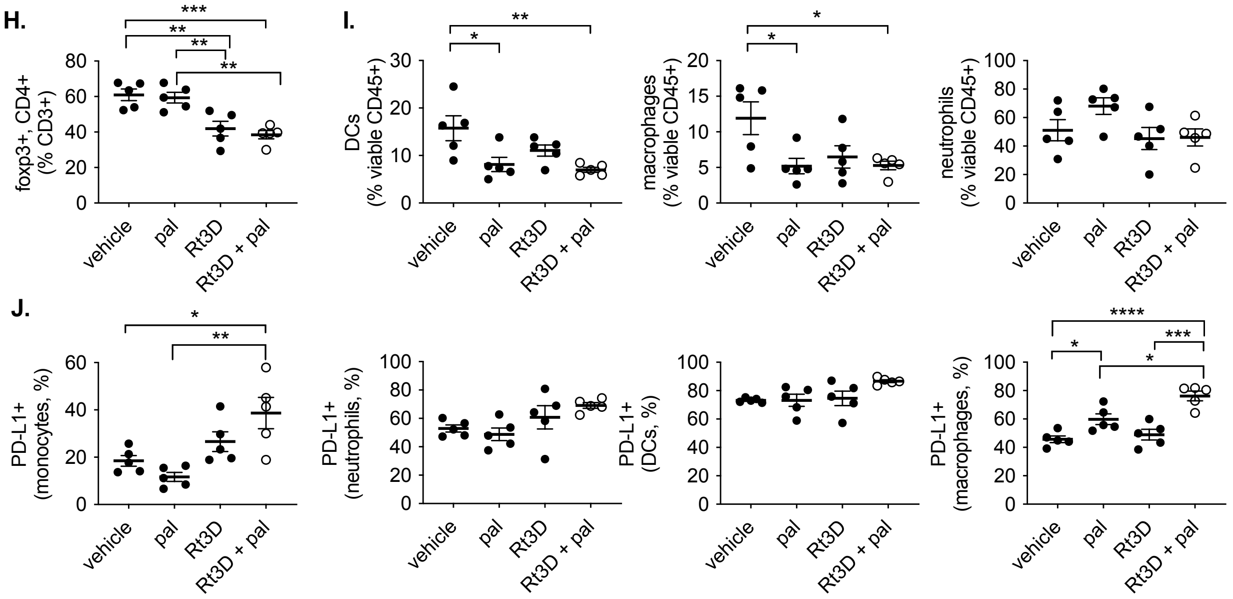


**Supplementary figure 2. A,** Proteomic analysis of A375 cells treated with Rt3D (0.1-1) in combination with palbociclib (1 µM) reveal upregulated (red) and downregulated (green) proteins categorised under the GO term 'immune response' and **B,** 'IFN regulation and response' at 48 hours. **C,** Heat-map of RNA sequencing showing upregulated (red) and downregulated (blue) HLA-related genes for A375 cells treated with Rt3D plus palbociclib at 48 hours. **D,** Expression of HLA-A,B,C (class I) and HLA-DR,DP,DQ (class II) in A375 cells treated with Rt3D +/- palbociclib by FACS analysis showing 2 independent experiments. **E,** Expression of HLA class I and II in A375 cells treated with Rt3D plus palbociclib +/- the JAK/STAT inhibitor, ruxilitinib (RUX). **F,** Proteomic analysis of A375 cells treated with Rt3D (0.1-1) +/- palbociclib (1 µM) reveal upregulated (red) and downregulated (green) proteins (24 hours, above, 48 hours, below). **G,** FACS plots of macrophages stained with CD11b-FITC co-cultured with treated A375 tumour cells stained with phrodo showing double-stained (engulfed) cells, representative of 3 biological repeats (± SEM, *n* = 3). **H,** C57BL/6 mice bearing MOC1 tumours were treated with palbociclib (100 mg/kg daily by oral gavage). After 3 doses of palbociclib, tumours received a single injection of 5 x $10^6$ pfu Rt3D and harvested 7 days after a single injection of 5 × $10^6$ pfu Rt3D or sham injection and stained with extra- and intra-cellular antibodies to profile the immune infiltrate. Data show proportions (through percentage analysis) of foxp3+ CD4+ cells, as a percentage from the CD3+ population gated from viable cells. **I,** Proportions show (through percentage analysis) of CD11c+ MHCII+ stained cells (labelled dendritic cells, DCs), CD11b+ F480+ cells (labelled macrophages) and CD11b+ Ly6G+ cells (labelled neutrophils), all gated from the viable CD45+ gated population for each treatment arm. **J,** PD-L1 expression on CD45+CD11b+Ly6C+ cells (monocytes), neutrophils, DCs and macrophages as described, for each treatment arm. A one-way ANOVA was used with the p value corrected for multiple comparisons against the Rt3D + pal group. N=6 animals per group.
