## Supplemental figure 3 for "CDK4/6 inhibition and dsRNA sensor agonism co-operate to enhance anti-cancer effects through ER stress and immune modulation of tumour cells"

**Supplementry figure 3.**

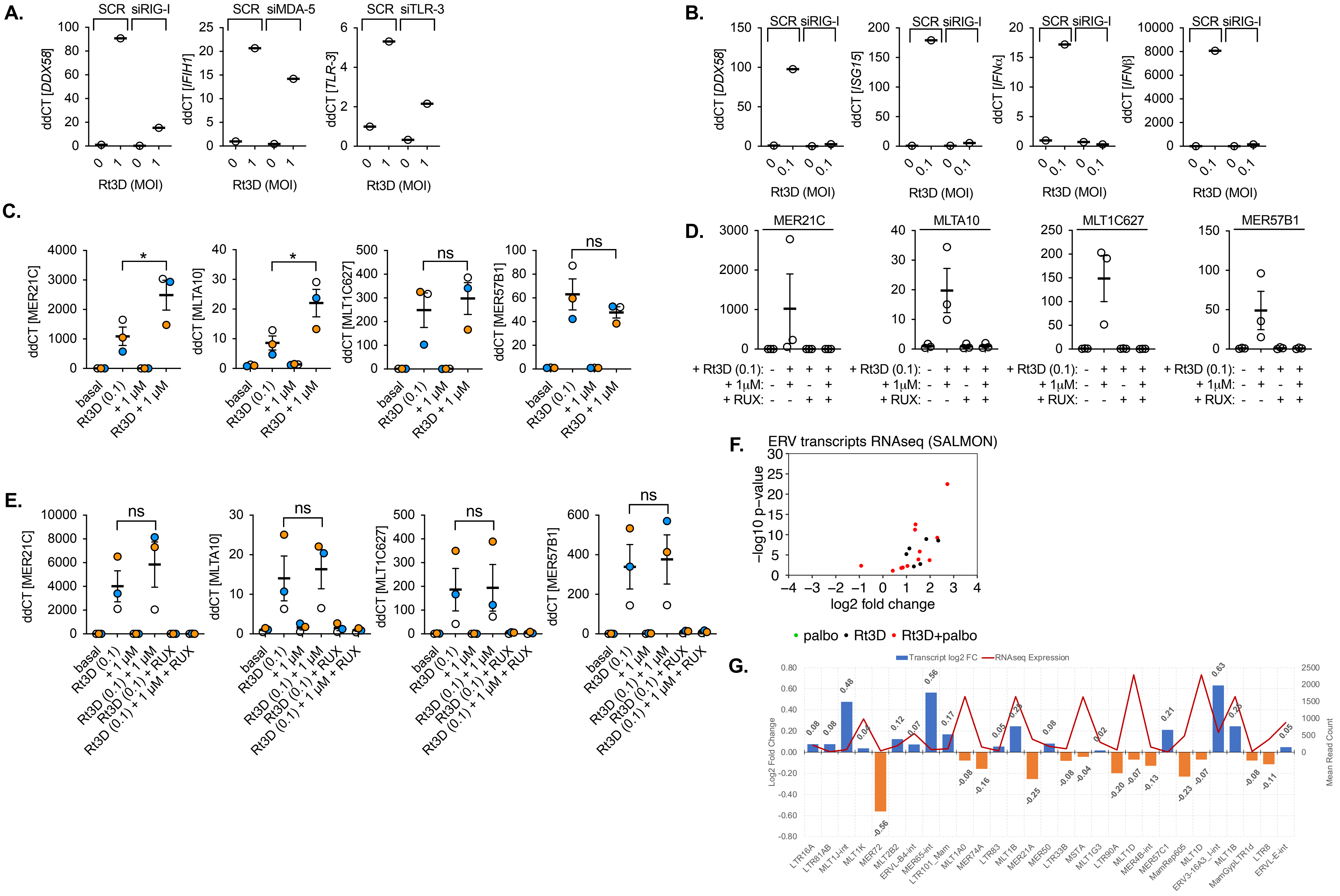

**H.**

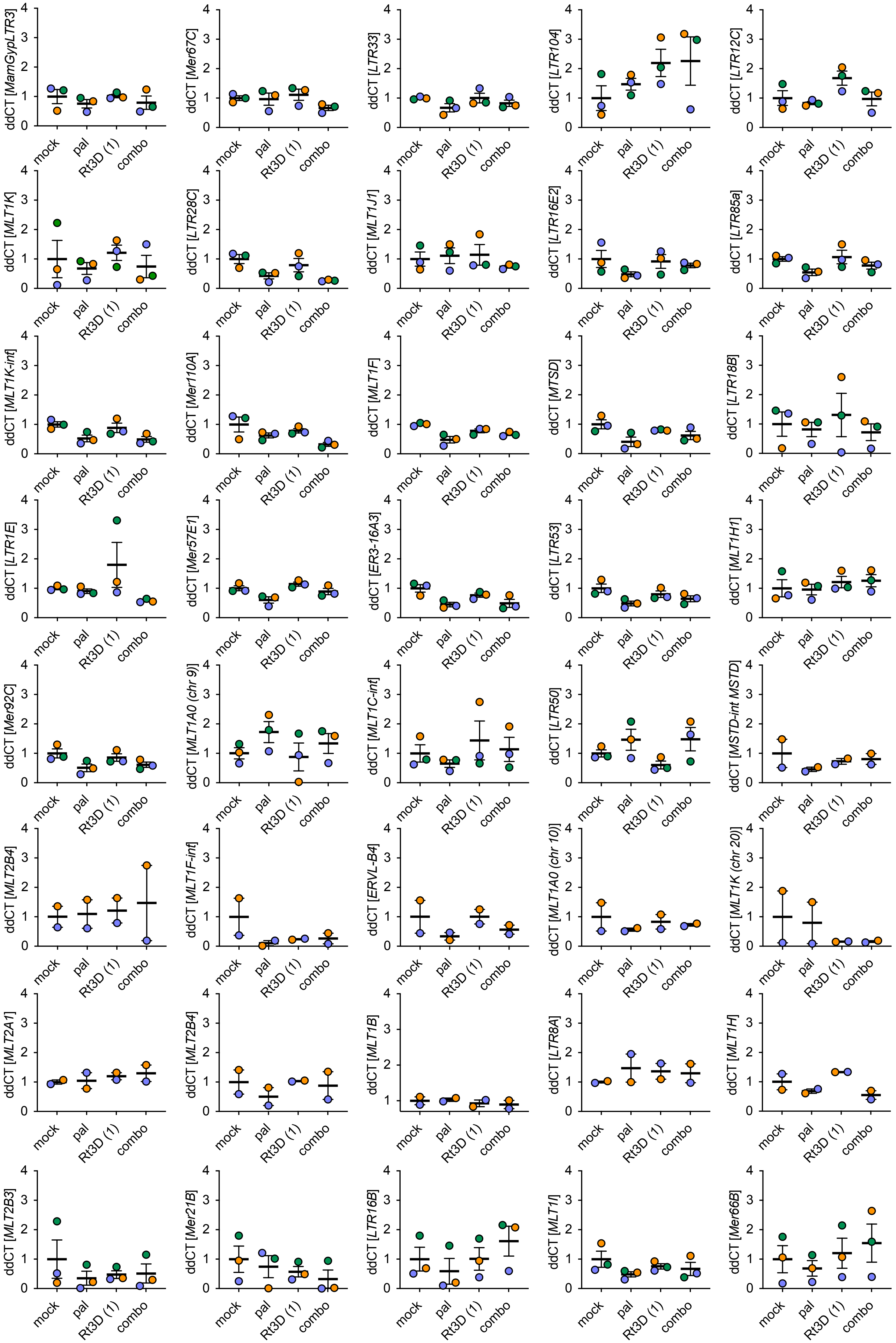

ddCT [MamGypLTR3]
ddCT [Mer67C]
ddCT [LTR33]
ddCT [LTR104]
ddCT [LTR12C]
ddCT [MLT1K]
ddCT [LTR28C]
ddCT [MLT1J1]
ddCT [LTR16E2]
ddCT [LTR85a]
ddCT [MLT1K-int]
ddCT [Mer110A]
ddCT [MLT1F]
ddCT [MTSD]
ddCT [LTR18B]
ddCT [LTR1E]
ddCT [Mer57E1]
ddCT [ER3-16A3]
ddCT [LTR53]
ddCT [MLT1H1]
ddCT [Mer92C]
ddCT [MLT1A0 (chr 9)]
ddCT [MLT1C-int]
ddCT [LTR50]
ddCT [MSTD-int MSTD]
ddCT [MLT2B4]
ddCT [MLT1F-int]
ddCT [ERVL-B4]
ddCT [MLT1A0 (chr 10)]
ddCT [MLT1K (chr 20)]
ddCT [MLT2A1]
ddCT [MLT2B4]
ddCT [MLT1B]
ddCT [LTR8A]
ddCT [MLT1H]
ddCT [MLT2B3]
ddCT [Mer21B]
ddCT [LTR16B]
ddCT [MLT1I]
ddCT [Mer66B]
mock
pal
Rt3D (1)
combo

**Supplementary figure 3.** **A,** RT-qPCR of RIG-I (DDX58), MDA-5 (IFIH1) or TLR3 at 48 hours following Rt3D infection (MOI 1) in A375, either transfected with scrambled (SCR) or siRNA knockdown of relevant targets. **B,** RT-qPCR of RIG-I (*DDX58*), *ISG15*, *IFN*, and *IFN*, at 48 hours following Rt3D infection (MOI 0.1) in A375, either transfected with siRNA against RIG-I or scrambled control (SCR). **C,** RT-qPCR of the ERVs *MER21C*, *MLTA10, MLT1C627* or *MER57B1*, in A375 cells treated with Rt3D (MOI 0.1) +/- palbociclib at 48 hours (± SEM, *n* = 3). **D,** *MER21C*, *MLTA10, MLT1C627* or *MER57B1* expression in A375 cells treated with Rt3D (MOI 0.1) and/or palbociclib (1 µM) +/- the JAK/STAT inhibitor, ruxolitinib (RUX, 1 µM) at 48 hours (± SEM, *n* = 3). **E,** RT-qPCR of *MER21C*, *MLTA10, MLT1C627* or *MER57B1* in *RB*-null A2058 cells treated with Rt3D +/- palbociclib (1 µM), +/- ruxolitinib (RUX), at 48 hours (± SEM, *n* = 3). **F,** Volcano plot of RNA sequencing showing upregulated and downregulated ERVs for A375 cells treated with Rt3D plus palbociclib compared to indicated single agents at 48 hours using SalmonTE pipeline with FDR adjusted p-values. **G,** From RNAseq data, fold-change in ERV transcripts and mean abundance of ERV transcripts across the sample for the ERV peptides from captured HLA-I with Rt3D-palbociclib versus basal conditions (shown in **Fig. 3M**). **H,** RT-qPCR of ERVs captured from peptides from immunopeptidomic analysis of HLA-I on A375 cells treated with Rt3D +/- palbociclib (1 µM), at 48 hours (± SEM, *n* = 3).
