## Supplemental figure 4 for "CDK4/6 inhibition and dsRNA sensor agonism co-operate to enhance anti-cancer effects through ER stress and immune modulation of tumour cells"

### Supplementry figure 4.

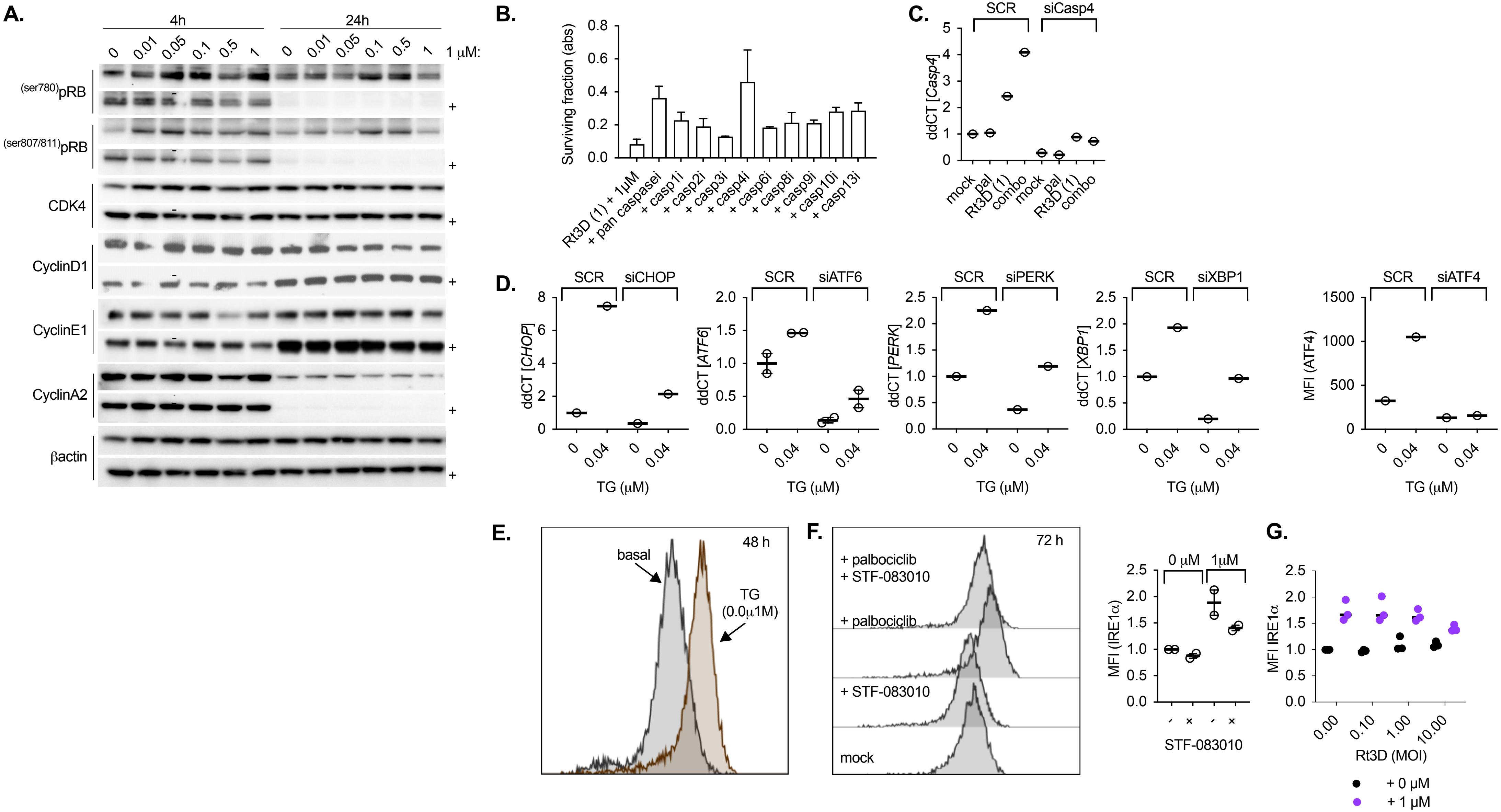


**Supplementary figure 4. A,** Western blot of cell cycle proteins treated with indicated doses of Rt3D with palbociclib (1 µM) in a A375 cells at 4 and 24 hours. **B,** A375 cells treated with combination therapy in the presence of either pan-caspase, or individual caspase inhibitors, and measured for cell survival by MTT assay at 72 hours. **C,** RT-qPCR of caspase 4 at 48 hours following palbociclib (1 µM), Rt3D (MOI 1), or the combination in A375, either transfected with scrambled (SCR) or with caspase 4 siRNA. **D,** Confirmation of on target effects of siRNA. RT-qPCR of CHOP, ATF6, PERK or XBP1 at 48 hours following thapsigargin (TG, 0.04 µM) in A375, either transfected with scrambled (SCR) or with the relevant siRNA. For confirmation of ATF4 siRNA, A375 cells containing an ATF4 reporter were treated with SCR or siATF4 and fluorescence measured at 72 hours. **E,** Validation of the IRE1 reporter cells. A375 cells containing an IRE1 endonuclease reporter was treated with thapsigargin (TG), to stimulate the UPR response. **F,** A375 cells containing an IRE1 endonuclease reporter were treated with palbociclib (1 µM) in combination with the IRE1 inhibitor (STF-083010). MFI is shown (right). **G,** A375 cells containing an IRE1 endonuclease reporter were treated with Rt3D (MOI 0.1-10) +/- palbociclib (1 µM) and subject to FACS analysis. Data show MFI, increased with palbociclib (± SEM, *n* = 3).
