## Supplemental figure 5 for "CDK4/6 inhibition and dsRNA sensor agonism co-operate to enhance anti-cancer effects through ER stress and immune modulation of tumour cells"

**Supplementry figure 5.**

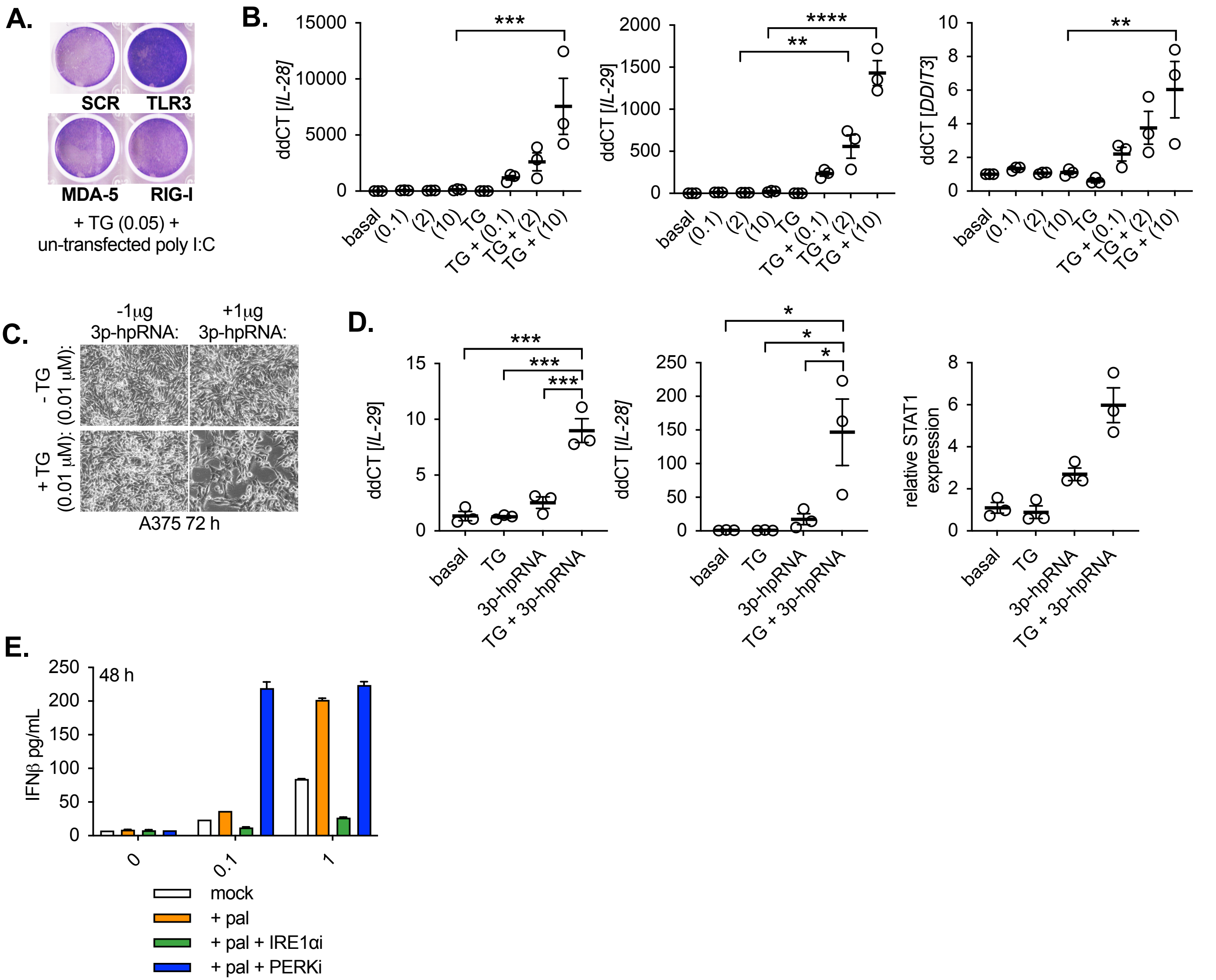


**Supplementary figure 5. A,** siRNA knock down of RNA sensors in A375 cells treated with untransfected poly I:C (10 µg/mL) plus TG (0.05 µM) showing cell viability by crystal violet. **B,** RT-qPCR of *IL-28*, *IL-29* and CHOP (*DDIT3*) in A375 cells treated with cells treated with transfected poly I:C at indicated doses, combined with thapsigargin (TG, 0.05 µM) at 48 hours. **C,** Pictomicrographs of A375 cells treated with the combination of 3p-hpRNA (1 µg/mL) plus TG (0.05 µM), at 72 hours. **D,** RT-qPCR of *IL-28*, *IL-29* or *STAT1* in A375 cells treated with cells treated with 3p-hpRNA, combined with thapsigargin at 72 hours (± SEM, $n$ = 3). **E,** Cells were treated with Rt3D (MOI 0.1, 1) +/- palbociclib, in combination with either the IRE1α inhibitor, or PERK inhibitor (GSK2656157, 2 µM) as indicated. IFN was measured in cell-free supernatant 48 hours later by ELISA. Data are representative of 2 independent experiments. A one-way ANOVA was used to compare IFN against samples treated with palbociclib for each dose of Rt3D as shown.
