## Supplemental table 1 for "CDK4/6 inhibition and dsRNA sensor agonism co-operate to enhance anti-cancer effects through ER stress and immune modulation of tumour cells"

| ERV | Forward (5'-3' | Reverse (5'-3') | BLASTn F | BLASTn R |
|---|---|---|---|---|
| *MLT1C49* | TATTGCCGTACTGTGGGCTG | TGGAACAGAGCCCTTCCTTG | TRIM22 | TRIM22 |
| *MER21C* | GGAGCTTCCTGATTGGCAGA | ATGTAGGGTGGCAAGCACTG | OAS2 | OAS2 |
| *MLTA10* | TCTCACAATCCTGGAGGCTG | GACCAAGAAGCAAGCCCTCA | DDX60L | DDX60L |
| *MLT1C627* | TGTGTCCTCCCCCTTCTCTT | GCCTGTGGATGTGCCCTTAT | OAS2 | OAS2 |
| *MER57B1* | CCTCCTGAGCCAGAGTAGGT | CCTCCTGAGCCAGAGTAGGT | OAS2 | OAS2 |

**Supplementary table 1. BLAST search for ERV primer sequences.**
BLAST search using primer sequences for 5 Rt3D-induced ERVs and the corresponding genes into which they align.
